# Genetic reassortment and diversification of host specificity have driven evolutionary trajectories of lineages of panzootic H5N1 influenza

**DOI:** 10.1101/2025.08.20.670882

**Authors:** William T. Harvey, Rute Maria Pinto, Maryn D. Brown, Lu Lu, Jessica L. Quantrill, Jiayun Yang, Nunticha Pankaew, Miranda Nel, James Baxter, Alexander M.P. Byrne, Darrell R. Kapczynski, Munir Iqbal, Joe James, Ashley C. Banyard, Ian Brown, Wendy Barclay, Thomas P. Peacock, Paul Digard, Samantha J. Lycett

## Abstract

Since 2021, subclade 2.3.4.4b A(H5N1) high pathogenicity avian influenza (HPAI) viruses have undergone changes in ecology and epidemiology, causing a panzootic of unprecedented scale in wild and domestic birds with spill-over infections and perceptible transmission in a range of mammalian species, raising concern over zoonotic potential. HPAI viruses readily exchange gene segments with low pathogenicity avian influenza viruses via reassortment, a mechanism that facilitates pronounced phenotypic change. Observations suggest changes in the seasonality and host range of panzootic viruses, however, data on the role of reassortment in determining such features are limited. Using phylodynamic approaches, we describe the emergence of the panzootic lineage and using a novel global genotype classification system we describe the subsequent emergence and global structuring of genotypes generated by reassortment. Focusing on evolutionary dynamics in Europe, we show reassortment has produced high fitness genotypes with enhanced capacity for transmission and further we show such advantages can be host-dependent, contrasting successful generalist genotypes with a specialist lineage (EA-2022-BB) adapted to birds of the order Charadriiformes. Experimental investigation of NS1-mediated shutoff indicates this Charadriiformes-specialist does not inhibit host cellular gene expression and hamper the defences of more typical hosts such as water- and land-fowl. We attribute this primarily to variation at position 127 of the NS1 protein. Our results emphasise that reassortment has driven phenotypic change, affected viral fitness, and caused diversification of host specificity and seasonality. Such factors should be considered in studies that seek to identify drivers of HPAI spread and map spillover risk. Additionally, relaxation of host specialisation, ecological diversification, and potential endemicity in atypical host populations present new reassortment opportunities that could result in further novel phenotypes.

## Introduction

Influenza A viruses (IAV), of the family Orthomyxoviridae, have segmented negative-sense single-stranded RNA genomes consisting of eight gene segments (1). Like other RNA viruses, IAVs exhibit a high mutation rate, typically accumulating between two to eight nucleotide substitutions per 1000 sites annually (2). Genome segmentation further facilitates the potential for rapid changes in viral characteristics through the process of genomic reassortment, which involves the exchange of gene segments between two (or more) viruses that coinfect the same host cell. The diversity of IAV in aquatic birds exceeds that of any other host group and is primarily supported by two highly divergent avian orders: Anseriformes (waterfowl, including ducks, geese and swans) and Charadriiformes (gulls, auks and shorebirds) (1,3). Reassortment is a common occurrence among avian IAVs, especially in wild bird populations, and is a major driver of virus diversity. Given the great diversity of IAVs in aquatic birds, reassortment between distantly related viruses can result in the rapid emergence of novel strains and phenotypes (4).

The ongoing avian influenza panzootic is caused by high pathogenicity avian influenza (HPAI) viruses belonging to the H5 Goose/Guangdong (Gs/Gd) lineage subclade 2.3.4.4b. HPAI viruses are differentiated from low pathogenicity avian influenza (LPAI) viruses by the presence of a multi-basic cleavage site in the haemagglutinin (HA) surface glycoprotein which enables intracellular processing of HA via ubiquitous furin-like proteases expanding tissue tropism and facilitating systemic disease (5,6). To date, HPAI viruses with multi-basic HA cleavage sites have been detected almost exclusively in the subtypes H5 and H7, and are estimated to have evolved independently from LPAI precursors on at least 39 occasions (4,7). In 1996, the HPAI H5N1 virus, A/Goose/Guangdong/1/96 was identified in commercial geese in Guangdong Province, China (8), and in the following year related viruses caused a significant outbreak of HPAI in poultry in Hong Kong and associated fatal human infections (9). In subsequent years, the evolution of the Gs/Gd lineage has involved frequent reassortment of “internal protein”-coding (*i*.*e*. non-glycoprotein) segments and diversification of the HA gene into ten major clades (10,11). Of these, clade 2 has descended several sub-clades notable for their overall frequency of detection, geographical spread, and spill-over to humans (4). Across most of the viral diversity in Gs/Gd clade 2 viruses, the H5 HA is typically paired with an N1 neuraminidase (NA). However, the subclade 2.3.4.4 is notable for undergoing frequent reassortment with LPAI strains since 2008; 2.3.4.4 HAs have circulated with a broad range of NA subtypes alongside N1 including N2, N3, N4, N5, N6 and N8. As such these viruses are collectively known as 2.3.4.4 H5Nx (4,11,12). The emergence of clade 2.3.4.4 H5Nx is linked to the increase in HPAI wild bird outbreaks outside Asia in the last decade; experimental studies have demonstrated a reduction in the pathogenicity of these viruses in wild ducks, facilitating global spread (4,12–14).

Since 2020, HPAI H5N1 viruses of subclade 2.3.4.4b emerged from, and replaced, previously circulating 2.3.4.4b H5N8 viruses. These H5N8 viruses had re-emerged in 2020 following a period of limited detections of 2.3.4.4b and an absence of a significant HPAI H5 wild bird epizootics since those caused by 2.3.4.4b H5N8 in 2016-2017 (15,16). Phylogenetic data indicate that the resurgent H5N8 originated from viruses related to those circulating in domestic birds in Egypt since 2018 (15,17,18). These viruses drove an epizootic in Europe in 2020/21 and reassorted with an LPAIV circulating in Europe resulting in the emergence of 2.3.4.4b H5N1 which, in addition to acquiring an N1 NA, acquired five internal gene segments (PB2, PB1, PA, NP and NS) (15,16). This 2.3.4.4b H5N1 lineage is estimated to have emerged in Europe or central Asia in late 2020 and has subsequently spread to Africa (15,16,19), Asia (15), North America (20,21), South America (22,23) and Antarctica (24). The ability of this H5N1 HA/NA combination to infect a broad range of species has resulted in extensive reassortment with non-notifiable LPAIVs, particularly in both Europe (16) and North America (20,25), to create a diverse array of genotypes with this conserved HA/NA pairing. The current situation represents an unprecedented level of global expansion for HPAI associated with changes in viral epidemiology and ecology that has coincided with devastating impacts on poultry and other farmed species. Further, infection of an unprecedented range of wild bird species has caused extensive mass mortality events globally, and the sheer number of dead wild birds has resulted in an increasing number of mammalian cases, often following scavenging activities, but also including multiple instances with evidence of mammal-to-mammal transmission (26–31).

Here we sought to evaluate the role of reassortment in the evolution and diversification of 2.3.4.4b H5N1 viruses following their emergence from H5N8 viruses. We present a system to classify all gene segments of subclade 2.3.4.4b H5Nx viruses and use these to classify genomes into profiles that reflect genotypes generated through genomic reassortment. We have used this global classification system to track segments and genotypes globally through time and to explore geographic structure. Tracking these genotypes using phylodynamic approaches, we address key questions in the evolution and epidemiology of panzootic HPAI viruses such as whether reassortment has generated genotypes with fitness advantages that increase the capacity of the virus to transmit within host populations or affected host specificity. To explore the mechanisms underpinning the observed ecological and evolutionary patterns, we experimentally characterised growth kinetics, polymerase activity and host shutoff of viruses representative of major reassortant lineages that have circulated in Europe. In an instance where there was evidence of differentiation in host specificity associated with reassortment, we explored the recent evolutionary history of the novel segments acquired by H5N1 viruses.

## Results

Following a period of low detections of 2.3.4.4b H5Nx across multiple years, a resurgence of H5N8 was seen in late 2019 and 2020, involving two genetically distinct H5N8 lineages that had diverged several years previously, according to the HA phylogeny (Fig 1a). Examination of time-resolved 2.3.4.4b HA phylogenies showed that in the late 2020 and early 2021, one of these H5N8 lineages underwent many changes in subtype through reassortment of the NA segment (Fig 1a, see Fig S1 for further detail). There is phylogenetic support for multiple reassortments from H5N8 to subtypes H5N1, H5N2, H5N5 and H5N6 and single transitions from H5N8 to H5N3 and H5N4. Additionally, there is strong support for reassortment from H5N5 to H5N3 in 2020 (Fig S1). Of the three H5N1 clades that emerged in late 2020, one has continued to circulate beyond 2022 and has caused a global panzootic showing transmission across Afro-Eurasia and in the Americas and Antarctica, as shown by the coloured bar alongside the phylogeny (Fig 1a). This panzootic H5N1 lineage, which comprises the upper portion of the phylogeny in Fig 1a, is estimated to have emerged around 20 August 2020 with a posterior distribution around this time of the most recent common ancestor (tMRCA) of 30 July to 22 September (95% highest posterior density (HPD)).

**Fig. 1.**
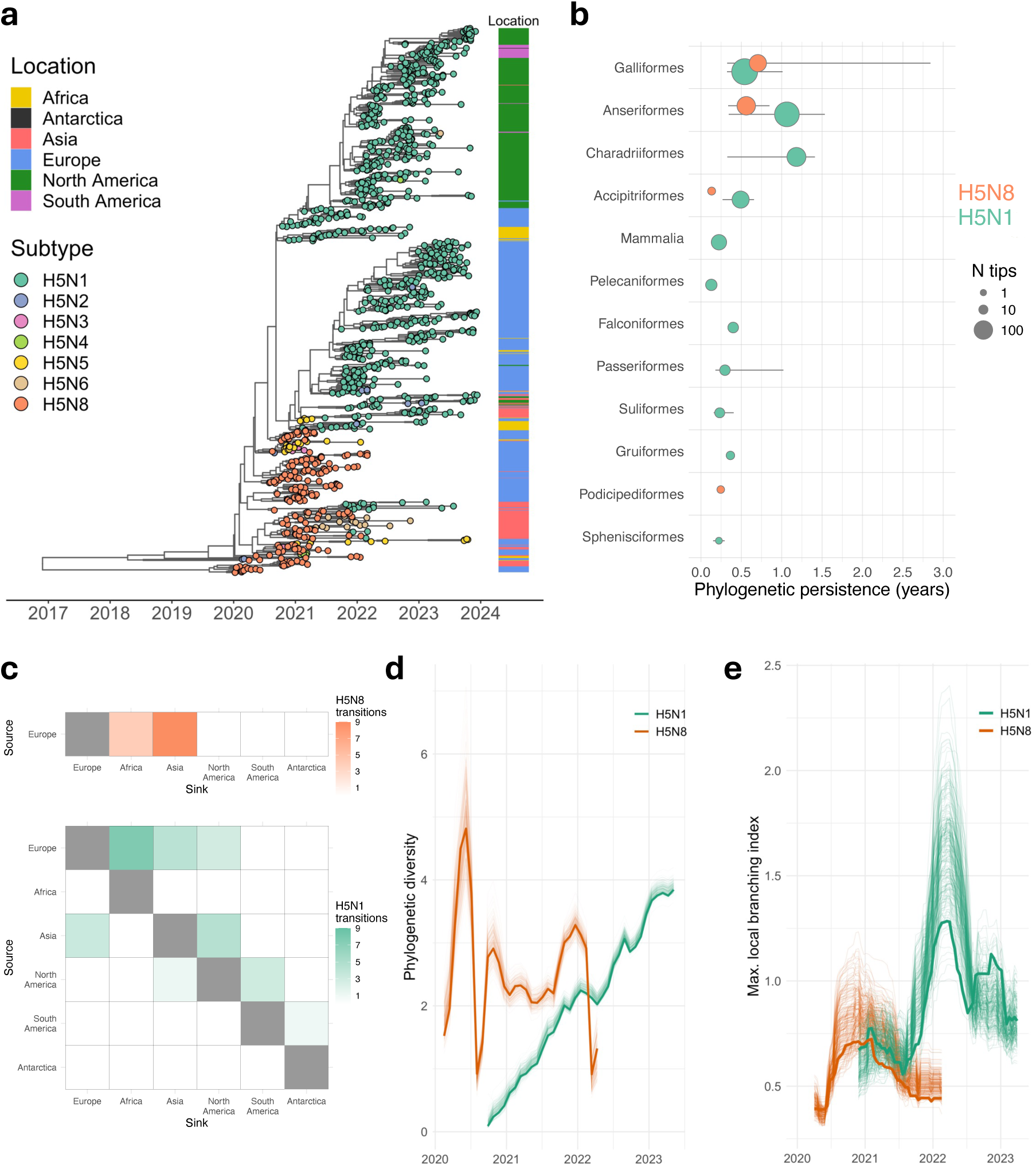
Phylogenetic, geographical and host association trends associated with the emergence of panzootic H5N1. **a**, Global time-scaled subclade 2.3.4.4b HA tree for geographically stratified sample of 1,073 viruses sampled between January 2020 and December 2023, subtype indicated by tip circle colour and sampling location at the level of continent shown in the coloured bar to the right. **b**, Duration of host-specific phylogenetic persistence for avian orders and for mammals. Circle position indicates median persistence with lines showing interquartile range and circle area indicates the number of tips. **c**, Phylogenetically robust counts of transitions between source and sink locations averaged across a posterior sample of 200 trees, with subtype and location reconstructed as discrete traits. **d**, HA phylogenetic diversity per subtype calculated as the mean pairwise patristic distance among H5N1 and H5N8 viruses sampled within a quarterly sliding window. **e**, Fitness of each subtype approximated from shape of tree with internal nodes labelled by subtype and local branching index through time calculated per-node using a sliding window. For **d** and **e**, thick and thin lines are based on the maximum clade credibility tree and a posterior sample of 200 trees, respectively.

To investigate potential differences in the host types in which viruses of different subtypes were transmitted and evolved within, phylogenetic persistence by host type was calculated. Phylogenetic persistence increases when a trait, such as host type, reconstructed at internal nodes of the phylogeny, shows a tendency to occur at deeper nodes of the phylogeny (32). For a host type with several detections over time, low values of phylogenetic persistence indicate a phylogeny consistent with repeated spillovers into this host with limited or shorter transmission chains thereafter. In contrast, higher values of phylogenetic persistence are consistent with longer chains of transmission within that host type. In Fig 1b, phylogenetic persistence for different host types is shown for H5N1 and H5N8, while the number of detections is indicated by circle area. Among the most frequently encountered host types, there are stark differences in persistence estimates between subtypes and there are several avian orders for which there is a measurement for H5N1 but not H5N8, a sign that those host types were not sampled with H5N8 or that they occurred only as isolated tips in the phylogeny. A further dramatic difference was the inference of high phylogenetic persistence of H5N1 in Charadriiformes, exceeding one year. In contrast, there was no evidence for persistence of H5N8 in Charadriiformes. Indeed, the average phylogenetic persistence of H5N1 in Charadriiformes (median = 1.2y, interquartile range (IQR) 0.3-1.4y) was slightly higher than the corresponding value in Anseriformes (median = 1.1, IQR 0.3-1.5y), which, as the presumed wild bird reservoir for 2.3.4.4b H5 viruses, were expected to exhibit high persistence estimates. Notably for H5N8, the highest persistence estimate is for Galliformes (median = 0.7y, IQR 0.3-2.8y) which is consistent with the assumed period of endemicity in poultry of subclade 2.3.4.4b H5N8 that preceded their resurgence in 2019 (17). Here, estimates of phylogenetic persistence of H5N1 in mammals exist though remain low as this analysis covering the period of transition in subtype dominance precedes the establishment of the virus in dairy cattle in 2024, though it did include sequences from farmed mink and from sea lions, for which transmission is proposed. Overall, these phylogenetic inferences are consistent with either a relaxation of host specificity of H5N1, an increase in spillover events resulting in transmission chains simply because of elevated virus transmission in typical hosts, or both.

To explore differences in the global dispersal of H5N1 and H5N8 subclade 2.3.4.4b viruses during the period of transition in subtype dominance summarised in Fig 1, both subtype and continent were reconstructed as discrete traits across the phylogeny. Phylogenetically robust transitions, that appeared one or more times per tree averaging across a posterior sample of trees, were identified and are represented in heatmaps in Fig 1c. Spread of H5N8 was restricted geographically to Eurasia and Africa, with several reconstructed movements from Europe to both Asia and Africa (Fig 1c, upper panel). In contrast, H5N1 shows a greater diversity of intercontinental spread. Among these data, Europe is again inferred as the primary source for intercontinental movements of H5N1 with phylogenetically supported exports to Asia, Africa and North America (Fig 1c, lower panel). From Asia, there is support for exports to both Europe and North America. Of the various imports to North America, a lineage seeded by import from Europe via the North Atlantic route was initially far larger (comprising the upper section of the phylogeny in Fig 1a) and was subsequently exported on multiple occasions to South America, and from there to Antarctica. The scale of epizootic seeded by the Pacific route of transmission to North America was initially restricted in size, though it continued at lower levels for around two years before expanding in birds and causing notable infections in swine, cattle, and in humans in late 2024 and early 2025 (33). The geographic expansion of H5N1 was accompanied by increasing phylogenetic diversity (Fig 1d). In early 2020, H5N8 HA phylogenetic diversity was initially high due to the aforementioned presence of two divergent lineages, though this reduced dramatically when detection of one of these lineages largely ceased. It then remained steady through 2021 before collapsing in early 2022 as 2.3.4.4b H5N8 detections waned. Following emergence in late 2020, phylogenetic diversity of HA in H5N1 viruses increased steadily and exceeded that of the H5N8 viruses by the end of 2022.

To assess the epidemiological fitness of subtypes through time, the local branching index (LBI) was calculated from the shape of the phylogeny using a sliding window approach (Fig 1e). In addition to adaptive and maladaptive mutations in HA, these LBI calculations are also affected by other factors that impact virus transmission and thus the branching structure of the HA phylogeny. Most relevantly, it is affected by i) contributions of other gene segments to overall viral fitness and ii) environmental factors such as seasonal changes in host population structure. Upon emerging, the LBI of H5N1 rapidly exceeded that of circulating H5N8 viruses indicating a fitness advantage (Fig 1e). The effect of seasonality on transmission is also apparent with both subtypes showing peaks in LBI during winters of the northern hemisphere where most detections occurred. However, the height of winter peaks for H5N1 and indeed its summer trough in 2022 exceeded the winter 2020/21 H5N8 peak, emphasising the high fitness of H5N1 viruses circulating across this period. There is growing evidence that the increased fitness of subclade 2.3.4.4b H5N1 relative to the H5N8 precursor, apparent here in the shape of the HA tree, was conferred principally by other segments acquired through reassortment rather than by adaptive mutations in HA. As the panzootic H5N1 lineage has spread into new geographical regions of the world and circulated in populations of different host types, novel opportunities for reassortment with LPAI viruses circulating in different host populations have arisen, raising the potential for the emergence of phenotypically distinct reassortants.

### Global reassortment patterns

To explore intra-subtype reassortment events involving 2.3.4.4b H5Nx, genetic clusters were first defined independently for each gene segment from phylogenies generated from alignments of unique nucleotide sequences. Thus, for each genome, eight phylogenetic cluster identifiers were computed, and each combination of these identifiers was considered as a genomic profile. In Fig 2a, a time-scaled HA phylogenetic tree is shown for a geographically stratified sample of H5N1 viruses belonging to the panzootic lineage sampled between late 2020 and late 2024 and a small number of closely related H5N8 viruses. Alongside a schematic diagram illustrates the results of the phylogenetic clustering procedure with each column representing the results for a particular gene segment. The combination of colours across the set of eight columns, therefore, represents the reassortment profile. Reassortment events are visualised as colour changes in the segments acquired by subclade 2.3.4.4b viruses. For example, there are colour changes from light brown to pale green in six columns (representing PB2, PB1, PA, NP, NA and NS) associated with the emergence of panzootic H5N1 from its H5N8 precursor, while HA and M did not change. Genotype labels were assigned to profiles which indicate virus subtype, the year in which they were first detected, and an emergence, or ‘R’, number that denotes the sequence in which a genome was first sampled. Thus, the profile for the novel H5N1 genotype first detected in Europe in 2020, which subsequently descended the variety of genotypes associated with the global panzootic, was labelled H5N1/2020/R1. In Table 1, genotypes with more than 150 assigned genomes are shown alongside the continents in which they have been frequently sampled, the profile of segment identifiers and equivalent genotype designations made by established systems.

**Fig. 2.**
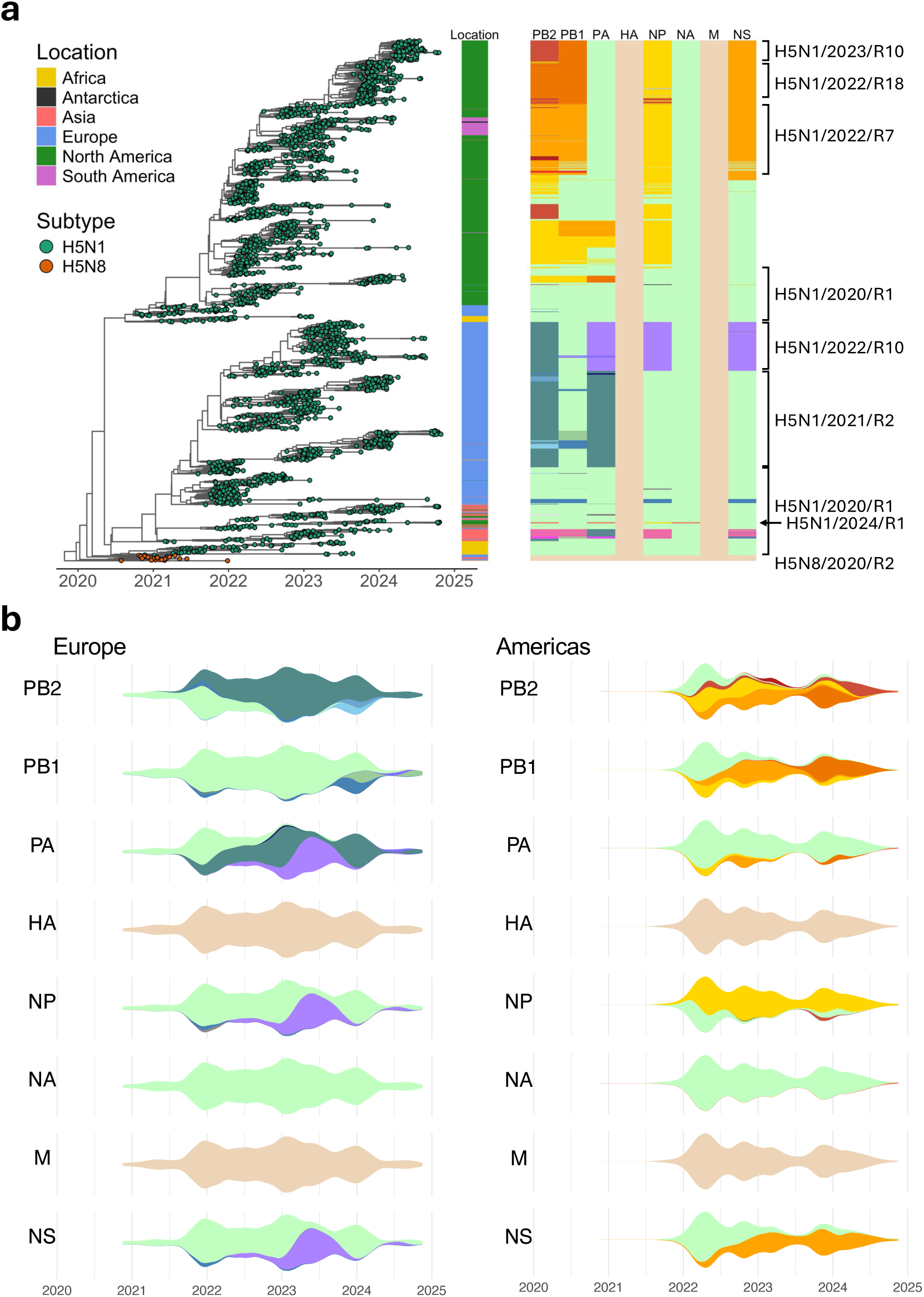
Reassortment and geographical compartmentalisation among panzootic H5N1 viruses. **a**, Global time-scaled subclade 2.3.4.4b HA tree for geographically stratified sample of 2,484 viruses sampled between August 2020 and October 2024, subtype indicated by tip circle colour and sampling location at the level of continent shown in a coloured bar. To the right, a schematic illustrates genetic cluster for each gene segment. Labels indicate the general areas of the schematic roughly corresponding to the six most represented H5N1 genotypes in this phylogeny. Additionally, the position of ancestral H5N8 viruses and early H5N1/2024/R1 (USDA D1.1) sequences are highlighted. **b**, Stream plots showing temporal changes in clusters for each gene segment, where height is proportional to the number of genomes represented in the schematic in **a** sampled from Europe (left) and from the Americas (right).

**Table 1.** Most frequently detected reassortant genotypes.

| Label | N | Principal geographies* | Profile | Comparable H5N1 genotype(s)<sup>†</sup> |  |
| --- | --- | --- | --- | --- | --- |
|  |  |  |  | Europe (EURL) | Americas (USDA) |
| H5N8/2020/R1 | 196 | Asia, Europe | 9_7_6_2_2_2_1_3 | - | - |
| H5N8/2020/R2 | 1,124 | Asia, Europe | 6_4_3_1_3_2_1_3 | - | - |
| H5N1/2020/R1 | 2,404 | Africa, Europe, N America | 3_1_1_1_2_1_1_1 | EA-2020-C | A1, A2, A3, A5 |
| H5N1/2021/R2 | 1,408 | Europe | 1_1_2_1_2_1_1_1 | EA-2021-AB | - |
| H5N1/2022/R1 | 322 | N America | 4_5_1_1_1_1_1_1 | - | B1.1 |
| H5N1/2022/R3 | 513 | N America | 3_1_1_1_1_1_1_1 | - | B4.1 |
| H5N1/2022/R4 | 864 | N America | 4_1_1_1_1_1_1_1 | - | B2.1 |
| H5N1/2022/R5 | 451 | N America | 4_5_5_1_1_1_1_1 | - | B1.2 |
| H5N1/2022/R6 | 199 | N America | 5_1_1_1_1_1_1_1 | - | B3.1 |
| H5N1/2022/R7 | 1,158 | N America, S America | 5_3_1_1_1_1_1_2 | - | B3.2 |
| H5N1/2022/R10 | 1,027 | Europe | 1_1_4_1_4_1_1_4 | EA-2022-BB | - |
| H5N1/2022/R12 | 202 | N America | 4_3_7_1_1_1_1_1 | - | B1.3 |
| H5N1/2022/R17 | 170 | Europe | 1_6_2_1_2_1_1_1 | EA-2022-CP | - |
| H5N1/2022/R18 | 644 | N America | 7_2_1_1_1_1_1_2 | - | B3.6 |
| H5N1/2023/R10 | 1,536 | N America | 3_2_1_1_1_1_1_2 | - | B3.13 |

\*Continents in which 10% or more genomes assigned to a reassortant genotype were sampled.
<sup>†</sup>Equivalent genotypes are listed if they match 5% or more genomes assigned to a genotype.

Following the emergence of the panzootic H5N1 lineage in 2020, there has been several exchanges of segments PB2, PB1, and PA which encode the tripartite RNA-dependent RNA polymerase, NP which encodes the viral nucleoprotein and NS which encodes the non-structural proteins NS1 and NS2/NEP (nuclear export protein). In contrast, exchanges of NA have occurred rarely and transiently, with the notable exception of the exchange of NA to a distinct N1 NA of North American origin in viruses labelled H5N1/2024/R1 in Fig 2a (equivalent to USDA D1.1). No exchanges of the M segment, which encodes the matrix proteins M1 and M2, were inferred. In the part of the tree representing the spread of subclade 2.3.4.4b viruses in Europe (sorted lower in Fig 2a), there are large blocks of teal and purple that correspond to exchanges of PB2, PA (twice), NP and NS. In the portion of the HA tree largely mapping to transmission in the Americas (sorted higher in Fig 2a), there are blocks of yellow, orange, and red in columns representing PB2, PB1, PA, NP and NS which indicate various acquisitions of these segments through reassortment with LPAI circulating in the Americas. Of all the segments first identified in profiles first observed in the Americas, we find no detections of these segments outside the Americas and Antarctica, except for a single virus of genotype H5N1/2022/R7 (USDA B3.2) sampled in Korea in 2022 (EPI_ISL_15647835). Notably, there is geographical segregation globally in the locations in which gene segments have been sampled, with acquired segments tending to have been sampled in either the Americas and Antarctica or in Afro-Eurasia. Given this geographic compartmentalisation and the absence of a globally dominant genotype, it is rational to examine the consequences of reassortment on transmission dynamics on a per-region basis.

### H5N1 reassortment dynamics in Europe

From late 2020 till the end of 2023, approximately 90% of subclade 2.3.4.4b H5N1 genomes sampled in Europe belonged to the reassortant genotypes H5N1/2020/R1 (EURL designation EA-2020-C), H5N1/2021/R2 (EA-2021-AB), and H5N1/2022/R10 (EA-2022-BB). These three genotypes are shown alongside other reassortants in a phylogeny of H5N1 viruses sampled in Europe between late 2020 and early 2024, where they are represented in blue, red, and green, respectively (Fig 3a, fully annotated version in Fig S3). The initial H5N1 genotype, H5N1/2020/R1, was ancestral to H5N1/2021/R2 which was first detected in 2021 and was inferred to have acquired novel segments PB2 and PA through reassortment. In turn, a virus assigned to the type H5N1/2021/R2 later reassorted resulting in the reassortant type H5N1/2022/R10 which was inferred to have undergone a further exchange of PA alongside acquiring novel NP and NS segments (Fig 2b). H5N1/2021/R2 emerged during the 2021/22 winter in Europe; its emergence coinciding with a seasonal peak in transmission of H5N1/2020/R1 as estimated from the branching pattern of the HA phylogeny (Fig 3b). Still, H5N1/2021/R2 appears to have an LBI indicative of increased fitness as it rapidly acquired a transmission advantage relative to H5N1/2020/R1 and LBI estimates remain higher for this genotype moving towards mid-2022. As H5N1/2021/R2 experienced a seasonal trough in transmission in the summer of 2022, the reassortant H5N1/2022/R10 emerged with initial transmission success apparent as a peak in LBI in mid-2022. In a multitype-tree birth-death model analysis of the emergence of H5N1/2022/R10, the effective reproductive number (*R_e_*) for this reassortant was estimated as quickly being greater than that for H5N1/2021/R2 and remaining so for the summer of 2022 (Fig S5). This represents the start of a period of asynchronous peaks and troughs in fitness estimates for the two reassortant types (Fig 3b). H5N1/2021/R2 recovered to a second peak in winter 2022/23, declining to a trough in mid-2023 before showing signs of recovery to a lesser degree in late 2023. Over the same period, H5N1/2022/R10 shows a decrease from its initial summer peak in LBI moving towards a trough early in winter of 2022/23, followed first by a large peak in mid-2023, and then by a clear decrease in late 2023. Therefore, H5N1/2021/R2 shows a pattern typical of H5s peaking in the northern hemisphere winter whereas H5N1/2022/R10 appears atypical, showing greater transmission capability in spring-summer.

**Fig. 3.**
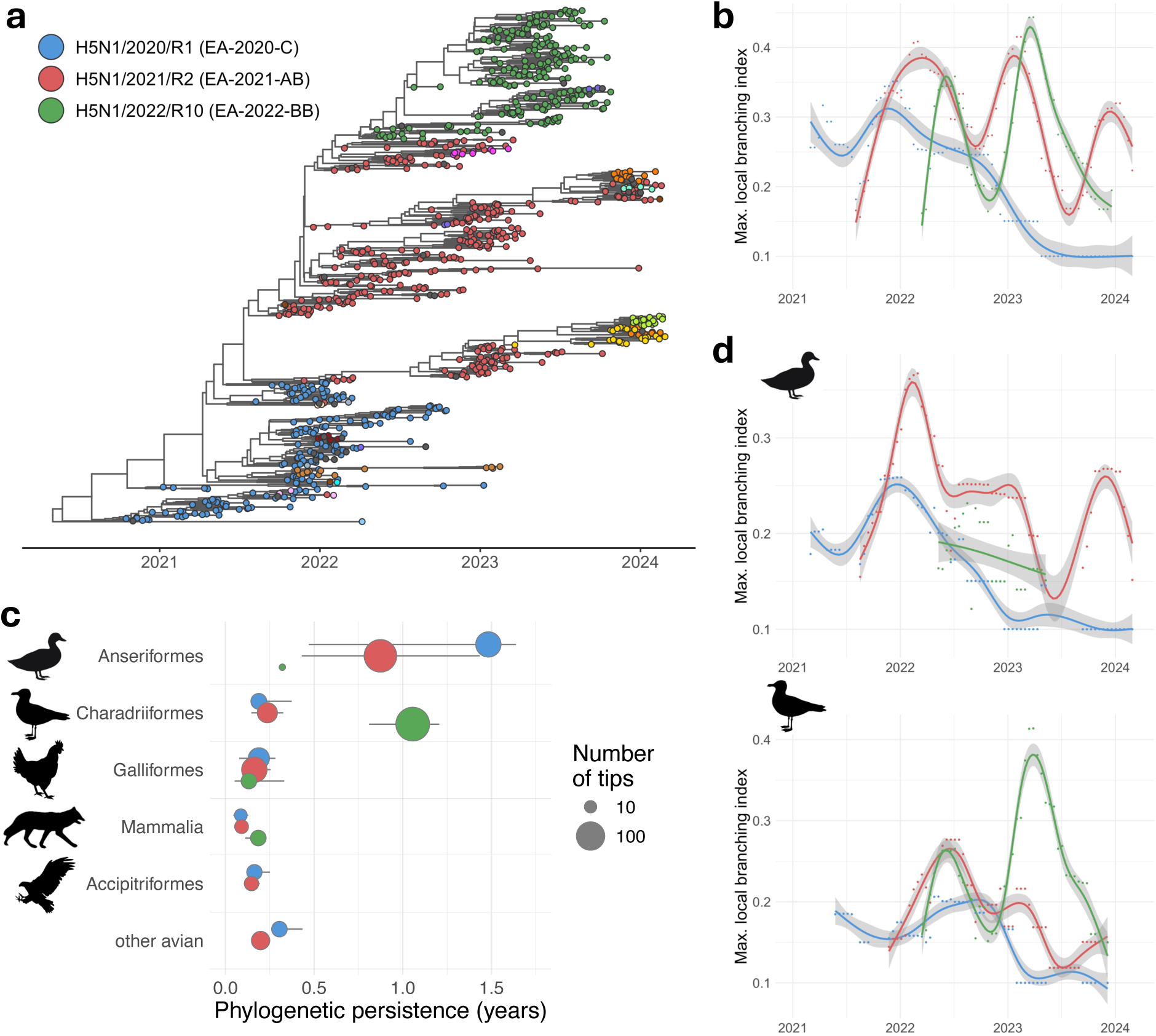
Fitness, seasonality and host preference vary between H5N1 reassortants. **a**, Time-scaled subclade 2.3.4.4b HA tree for geographically stratified sample of 875 H5N1 viruses sampled in Europe between October 2020 and February 2024, reassortant indicated by tip circle colour. **b**, Fitness of each major reassortant, across all hosts, approximated from shape of tree with local branching index calculated per-node using a temporal sliding window. Points show maximum values per-reassortant for each time window and curves are fitted to these using generalized additive models and visualised with 0.95 confidence intervals. **c**, Host-specific phylogenetic persistence summarised for the three genotypes H5N1/2020/R1, H5N1/2021/R2 and H5N1/2022/R10. Point position indicates median persistence and size indicates the number of tips with lines representing interquartile range. Calculated at the level of order for birds and for the class Mammalia and plotted for the five most frequently sampled host groups with remaining avian orders grouped under *other avian*. **d**, Host–specific tree-shape based fitness calculated using subtrees comprised of viruses sampled from Anseriformes (top) or Charadriifomes (bottom).

To investigate differences in host specificity that may have resulted from the reassortment events that have determined fitness dynamics in Europe, we quantified host-specific phylogenetic persistence for each reassortant genotype in different avian orders and for mammals, by reassortant type. Among the starkest differences in the most frequently sampled hosts (Fig 3c), we found that the phylogenetic persistence of H5N1/2022/R10 was high in Charadriiformes (median = 1.1y, IQR 0.8-1.2y) and relatively low in Anseriformes (median = 0.32 across three measures). While the other major reassortants were also frequently detected in Charadriiformes (21 H5N1/2020/R1 and 41 H5N1/2021/R2 in the down-sampled dataset compared with 75 and 141 genomes from Anseriformes, respectively), persistence estimates were lower indicating shorter, on average, transmission chains in Charadriiformes. After down-sampling, both H5N1/2020/R1 and H5N1/2021/R2 were most frequently detected in and showed highest persistence in Anseriformes (median 1.5y IQR 0.5-1.6y and median 0.9y IQR 0.4-1.4y, respectively). The moderate persistence of H5N1/2020/R1 and H5N1/2021/R2 in the group ‘other avian’, further indicates the generalist phenotype of the emergent panzootic clade. For H5N1/2020/R1, this was largely due to persistence of H5N1/2020/R1 in the northern gannet (*Morus bassanus*), a colony-nesting sea bird belonging to the order Suliformes with persistence estimated to have been second only to Anseriformes for this genotype. For H5N1/2020/R2, this was due to some inferred persistence in birds of the order Pelecaniformes such as the grey heron (*Ardea cinerea*). Persistence profiles and detection frequency for H5N1/2020/R1 and H5N1/2021/R2 in different host types are broadly similar indicating both viruses to be generalists, being able to infect and be shed and primarily maintained through transmission in Anseriformes. In contrast, H5N1/2022/R10 has a very distinctive persistence profile, with high persistence and greater specificity to Charadriiformes and relatively few detections in Anseriformes. Phylogenetic persistence estimates and detections in mammals, after down sampling to account for differences in sampling intensity, were broadly similar for each of these three genotypes, suggesting an absence of evidence of differentiation in potential for infection and transmission in mammals associated with these reassortment events.

Differences in host specificity have the potential to influence fitness dynamics shown in Fig 3b. To explore this explicitly, we investigated host-specific instances of the LBI calculated using the shape of subtrees comprised of branches ancestral to viruses sampled in the two host types inferred to show highest persistence in either of the principal reassortants, Anseriformes and Charadriiformes (Fig 3d). These host-specific instances were compared with the trends in multi-host LBI (Fig 3b). The trend in Charadriiformes-specific LBI for H5N1/2022/R10 closely follows the multi-host estimates for H5N1/2022/R10, confirming the close association of this reassortant to this host group. For H5N1/2020/R1 and H5N1/2021/R2, neither host-specific LBI trend (Fig 3d) is as well matched to their multi-host pattern (Fig 3b), though in both cases the Anseriformes-specific trend is closer. For H5N1/2021/R2, the peak in multi-host LBI observed in winter 2022/23 (Fig 3c) is not recapitulated in the anseriform-specific LBI. This suggests that the transmission of this reassortant in this period involves transmission in other host types in addition to Anseriformes. H5N1/2021/R2 also shows a peak in Charadriiformes-specific LBI in the summer of 2022 when its multi-host measure is falling suggesting a significant degree of circulation of this reassortant in charadriiforms, though coupled with low persistence (Fig 3c) indicative of short transmission chains within this host type. Together these results suggest H5N1/2020/R1 and H5N1/2021/R2 to be somewhat more generalist in their host specificity in comparison with H5N1/2022/R10 which is a Charadriiformes specialist.

### Major European genotypes differ in viral growth kinetics, polymerase activity and shutoff of host gene expression

To assess whether observed ecological differences between the three major H5N1 reassortant lineages could be replicated *in vitro,* representative viruses from each reassortant type were selected (Table 2): H5N1/2020/R1 (EA-2020-C) was represented by A/chicken/England/053052/2021 (AIV07), H5N1/2021/R2 (EA-2021-AB) by A/chicken/Scotland/054477/2021 (AIV09), and H5N1/2022/R10 (EA-2022-BB) by A/chicken/England/085598/2022 (AIV48). These three genotypes contain very similar HA and NA genes, with major differences between one another mapping to the internal gene segments. Therefore, virus growth replication kinetics were investigated using reassortant viruses containing six internal gene segments (PB2, PB1, PA, NP, M and NS) from representative H5N1 viruses and segments encoding the surface glycoproteins from the laboratory-adapted strain, A/Puerto Rico/8/1934 (PR8). Virus replication was tested in a panel of avian cell lines derived from chickens (CLEC213 and DF-1), duck (CCL-141) and quail (QT35). A consistent pattern was observed for all tested cells with AIV07 consistently replicating to significantly higher titres beyond 36 hours post-infection, compared with both AIV09 and AIV48 which displayed very similar kinetics to one another (Fig 4a). While the success of H5N1/2021/R2 (EA-2021-AB) in non-charadriiform species is not reflected by the reduced virus propagation of AIV09 in chicken, duck and quail cells, the lower fitness of AIV48 in these selected cells is in accordance with the reduced spread of H5N1/2022/R10 (EA-2022-BB) in galliform or anseriform species observed in nature.

**Table 2.** Laboratory representative viruses.

| Genotype label | EURL equivalent | Laboratory representative |  |
| --- | --- | --- | --- |
|  |  | Isolate label | GISAID isolate ID |
| H5N1/2020/R1 | EA-2020-C | A/chicken/England/053052/2021 (AIV07) | EPI_ISL_9012457 |
| H5N1/2021/R2 | EA-2021-AB | A/chicken/Scotland/054477/2021 (AIV09) | EPI_ISL_9012696 |
| H5N1/2022/R10 | EA-2022-BB | A/chicken/England/085598/2022 (AIV48) | EPI_ISL_13782459 |

**Fig. 4.**
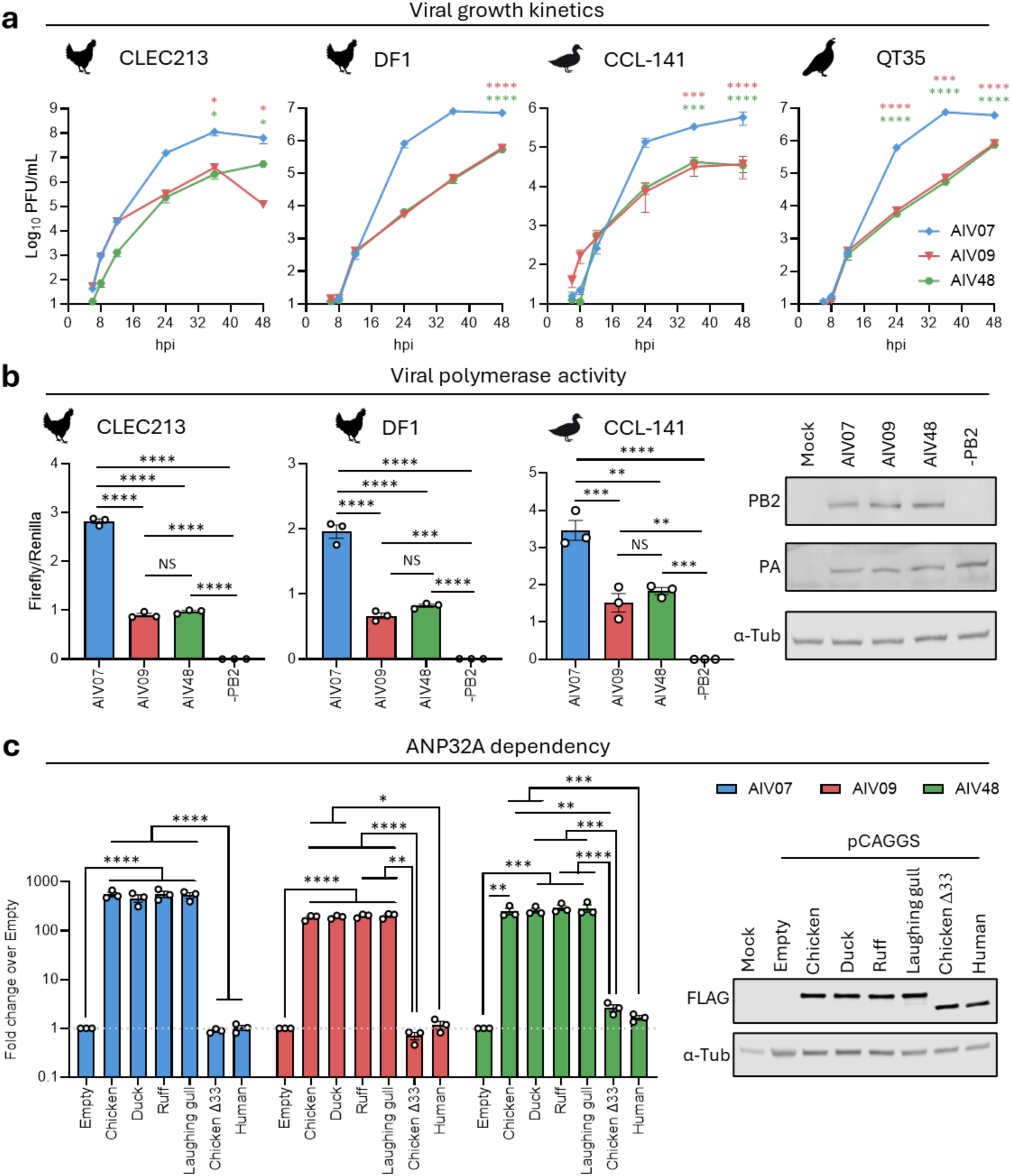
Reassorted viruses vary in replication kinetics and polymerase activity in cell culture but not ANP32 dependency. **a**, Replication in various cell types (CLEC213, DF1, CCL-141 and QT35) was assessed by infections conducted at multiplicity of infection (MOI) of 0.001 with 2:6 reassortments of AIV07, AIV09 and AIV48 with PR8 HA and NA. Supernatants were collected at various hours post-infection (hpi) and infectious viral titres determined by plaque assay. **b-c**, Polymerase activity was assessed by measuring luminescence 24 hours after transfecting cells with pCAGGs plasmids expressing PB2, PB1, PA and NP alongside a firefly luciferase-based vRNA-like reporter and a *Renilla* luciferase transfection control plasmid. A negative control was provided by transfection with PB2 pCAGGs replaced by empty pCAGGs (-PB2). Expression of PB2 and PA was detected by western blot. α-Tubulin was used as loading control. **c**, ANP32A-dependency was tested by the transfection of eHAPtKO cells with pCAGGs plasmids expressing PB2, PB1, PA, NP and various FLAG-tagged ANP32A homologues and mutants alongside a firefly luciferase-based vRNA-like reporter and a *Renilla* luciferase transfection control plasmid. Expression of FLAG-tagged ANP32A species was detected by western blot. α-Tubulin was used as loading control. For **a-c**, plotted data represent the mean ± SEM of technical repeats from three independent experiments. Statistical annotations (*p <0.05, **p <0.01, ***p <0.001, ****p <0.0001) are the result of one-way ANOVA comparisons with AIV07 (**a**) or as indicated (**b**-**c**).

Viral replication kinetics *in vitro* are the result of the complex contribution of the activities of several viral proteins encoded by multiple gene segments and the interaction between them. To investigate contributions of the gene segments involved in reassortment events separating these three lineages (PB2, PA, NP and NS segments), further experiments that measure functions of the proteins encoded by these segments were undertaken. Polymerase activity (primarily dependent on PB2, PB1, PA and NP proteins) was investigated using minireplicon reporter assays involving the transfection of CLEC213, DF1 and CCL-141 cells with plasmids expressing PB2, PB1, PA and NP from each of the representative viruses alongside a minigenome reporter expressing firefly luciferase. In Fig 4b, polymerase activity is shown as the firefly signal normalised to *Renilla* signal (firefly/*Renilla* ratios or relative luminescence units). Consistent with the virus replication experiments, AIV07 polymerase showed significantly higher polymerase activity than both AIV09 and AIV48 and no significant differences between the latter two were observed in any cell line, suggesting that differences in polymerase activity could be the major factor explaining the differing replication of these three viruses *in vitro*.

The influenza polymerase requires several host factors to efficiently replicate the virus genome (34). One well characterised host factor that can be responsible for interspecies differences in influenza virus susceptibility is the protein ANP32A (35). One possible explanation for the apparent differences in host range of the H5N1 genotypes prevalent in Europe could be differences in the compatibility between the viral polymerase complex and avian ANP32A proteins of different hosts. Therefore, we performed ANP32A reconstitution assays, by repeating minireplicon polymerase assays in a human cell line lacking pro-viral ANP32 proteins (eHAPtKO) (36), with complementation from ANP32 proteins from different avian species including representatives from the Galliformes (chicken), Anseriformes (duck) and Charadriiformes (ruff and gull) (Fig 4c). However, we saw no evidence of any differences in the compatibility between the different genotype polymerases and ANP32 proteins from different avian species, suggesting that intrinsic differences in virus polymerase and host ANP32A protein compatibility are unlikely to explain inferred differences in host specificity (Fig 4c).

To control host innate immune responses and virus-induced stimulation of interferon (IFN)-stimulated genes, influenza A viruses robustly antagonise type I IFN induction with antiviral properties. This is primary dependent on the functions of non-structural protein 1 (NS1) (37). Using reporter-based assays in a range of avian cells derived from both Galliformes and Anseriformes, we assessed the ability of NS segments from different reassortant types to counteract type 1 IFN induction following Polyinosinic:polycytidylic acid (Poly I:C) stimulation. However. NS segments from each of the representative viruses robustly antagonised type-I IFN induction and no variation between viruses was observed (Fig 5a). Moreover, influenza viruses can hamper host defences by inhibiting overall host cellular gene expression both at transcriptional and translational levels. This host shutoff is also mediated by NS1, and additionally by the PA-X protein, encoded by the PA segment and produced by programmed ribosomal frameshift (38). In experiments measuring PA-X-mediated shutoff, assays performed with the PA segment of these three viruses displayed broadly similar host shutoff (Fig 5b), with AIV48 showing marginally elevated shutoff in chicken cells. However, in experiments measuring NS1-mediated shutoff, a dramatically different phenotype was observed for AIV48 compared with either AIV07 or AIV09 (Fig 5c). Across the panel of avian cell lines, both AIV07 and AIV09 displayed consistently robust shutoff activity of around 80% or greater. In contrast, the NS1 segment of AIV48 showed no detectable shutoff activity in chicken and quail cells and only marginal shutoff (<20%) in duck cells.

**Fig. 5.**
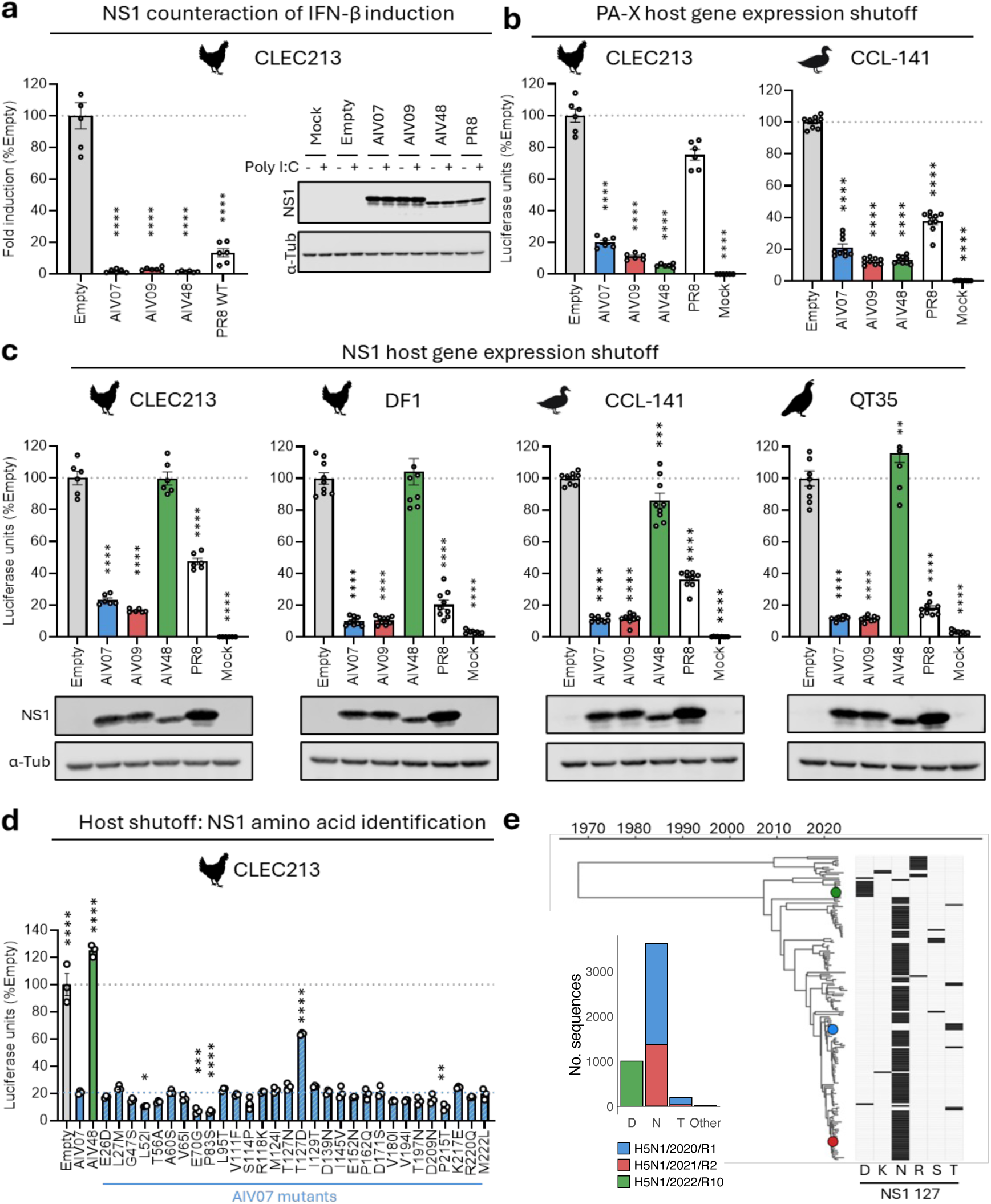
Reassorted viruses vary in NS1 mediated-host shutoff but not PA-X mediated-host shutoff or counteraction of IFN-β induction. **a**, CLEC213 and DF1 cells were co-transfected with pDUAL plasmids encoding segment 8 of the indicated viruses alongside a chicken IFN-β::firefly luciferase plasmid. Twenty-four hours post-transfection, cells were stimulated by Poly I:C transfection and luciferase levels were measured following a further 24h. PolyI:C driven luciferase induction was calculated and normalised to Empty plasmid. Expression of NS1 proteins in was assessed by western blot. α-tubulin was used as loading control. **b-c**, Increased PA-X (**b**) or NS1 (**c**) mediated host gene shutoff is represented by reduced luminescence measured at 48 hours post-transfection. Indicated cells were transfected with pDUAL plasmids encoding the PA (**b**) or NS (**c**) segment alongside a reporter plasmid that constitutively expresses *Renilla* luciferase. **d**, NS1-mediated host gene shutoff for AIV07 mutants with single amino acid substitutions present in AIV48 was assessed as in **c**. Dashed lines indicate mean luminescence measured for empty plasmid (grey) and AIV07 (blue) NS plasmid transfected cells. **e**, Time-scaled NS tree down-sampled to retain genetic diversity across subclade 2.3.4.4b H5 viruses sampled 2020-2024 with the positions of A/chicken/England/053052/2021 (AIV07), A/chicken/Scotland/054477/2021 (AIV09), and A/chicken/England/085598/2022 (AIV48) highlighted. Alongside, a schematic indicates amino acid at NS1 position 127. For **a**-**d**, the mean ± SEM of technical repeats from three independent experiments are plotted. Statistical annotations (*p <0.05, **p <0.01, ***p <0.001, ****p <0.0001) are the result of one-way ANOVA comparisons with values of empty (no NS1) plasmid-transfected cells (**a**-**c**) or with AIV07 (**d**).

To further investigate the molecular basis of the observed variation in NS1-mediated host shutoff, a comprehensive series of NS1 single amino acid mutants possessing AIV48 amino acids in an AIV07 background were generated. Characterisation of these mutants demonstrated that NS1 position 127 was the major determinant of the observed differences in shutoff with the shutoff activity of AIV07 reduced by greater than 50% when T127 was replaced by the D127 present in AIV48 (Fig 5d). While D127 present in AIV48 is typical of H5N1/2022/R10 viruses being present in around 97.8% of them (Fig 5e), T127 present in AIV07 was found in only a minority of H5N1/2020/R1 and H5N1/2021/R2 viruses (6.6% and 2.3%, respectively) whereas N127 was more commonly observed (93.1% and 97.2%, respectively). However, the substitution T127N was found not to alter NS1-mediated shutoff (Fig 5d) suggesting the observed differences in shutoff activity are likely generalisable across most viruses belonging to these genotypes. Together these results suggest that H5N1/2022/R10 viruses essentially lack NS1-mediated shutoff of host gene expression in cells derived from Galliformes and Anseriformes which could contribute to reduced fitness of this reassortant virus in these host types.

### Evolutionary origins of reassortant virus adapted to *Charadriiformes*

In the sprint to summer of 2022, H5N1/2022/R10 (EA-2022-BB) viruses emerged in Europe with novel PA, NP and NS segments acquired through reassortment. Given the particularly interesting change in host specificity displayed by these viruses, we investigated the evolutionary history of the novel genomic segments in greater detail. Early genomes of this genotype were sampled in countries bordering the North Sea, such as France, Belgium, and the Netherlands (16). Here, through analysis of the HA gene segment, we estimate an emergence time for this genotype as being around 3 April 2022 (95% HPD, 28 Feb-3 May) (Fig S6). It was noted that in the summer of 2022, while EA-2022-BB was the second most sequenced genotype in Europe, it was most frequently sampled in gulls (order Charadriiformes, family Laridae), particularly the European herring gull (*Larus argentatus*), while the most sampled species in 2023 was the black-headed gull (*Chroicocephalus ridibundas*) (16). For each of the reassorted segments (PA, NP and NS), sequences from non-H5N1 viruses most closely related to the earliest H5N1/2022/R10 sequence (A/gull/France/22p015977/2022 sampled on 11 May 2022, GISAID accession number: EPI_ISL_13519451) were identified using the basic local alignment search tool (BLAST) (39,40). Across these three segments, the 50 most closely related sequences from non-H5N1 viruses were from 73 viruses of subtypes H13N2 (n = 13 viruses), H13N6 (n = 21), H13N8 (n = 10), H13N9 (n = 1), and H16N3 (n = 28), with 70 of 73 viruses sampled from Charadriiforms (with waders and auks represented in addition to gulls and their allies) two from Anseriformes and an environmental sample (Table S1). All 50 NS sequences possessed D127 in NS1, which we associate with an absence of host shutoff in galliform and anseriform cells. H13 and H16 LPAI viruses are generally considered endemic in Charadriiformes and are hypothesised to have evolved endemically within birds of this order which are distantly related to Anseriformes (1,3,41).

For each of the segments, PA, NP and NS, time-scaled phylogenetic trees for 15 early H5Nx sequences and the 50 most closely related non-H5Nx sequences were generated (Fig 6a). In both the NP and NS trees, the sequences most closely related to those from H5N1/2022/R10 viruses come from an H13N8 virus sampled from a black-headed gull in Belgium in 2020 (A/Chroicocephalus ridibundus/Belgium/13464/2020, GISAID accession number: EPI_ISL_7622539). The PA from this virus is also closely related to the H5N1 reassortant that emerged in Europe, however for this gene segment there are closely related sequences sharing more recent common ancestors with the H5N1 viruses: an H13N6 virus sampled in South Korea and in H13N6/H13N2 viruses sampled in Delaware, USA. There are no genomes that possess an intermediate constellation of the reassorted segments that challenge the hypothesis that the PA, NP and NS segments were acquired by H5N1 in a single reassortment event with an H13 virus. The many switches in HA (and NA) subtypes across the phylogenies in Fig 6a and the discrepancies in topology emphasise the omnipresence of genomic reassortment among H13 and H16 LPAI viruses circulating in Charadriiformes.

**Fig. 6.**
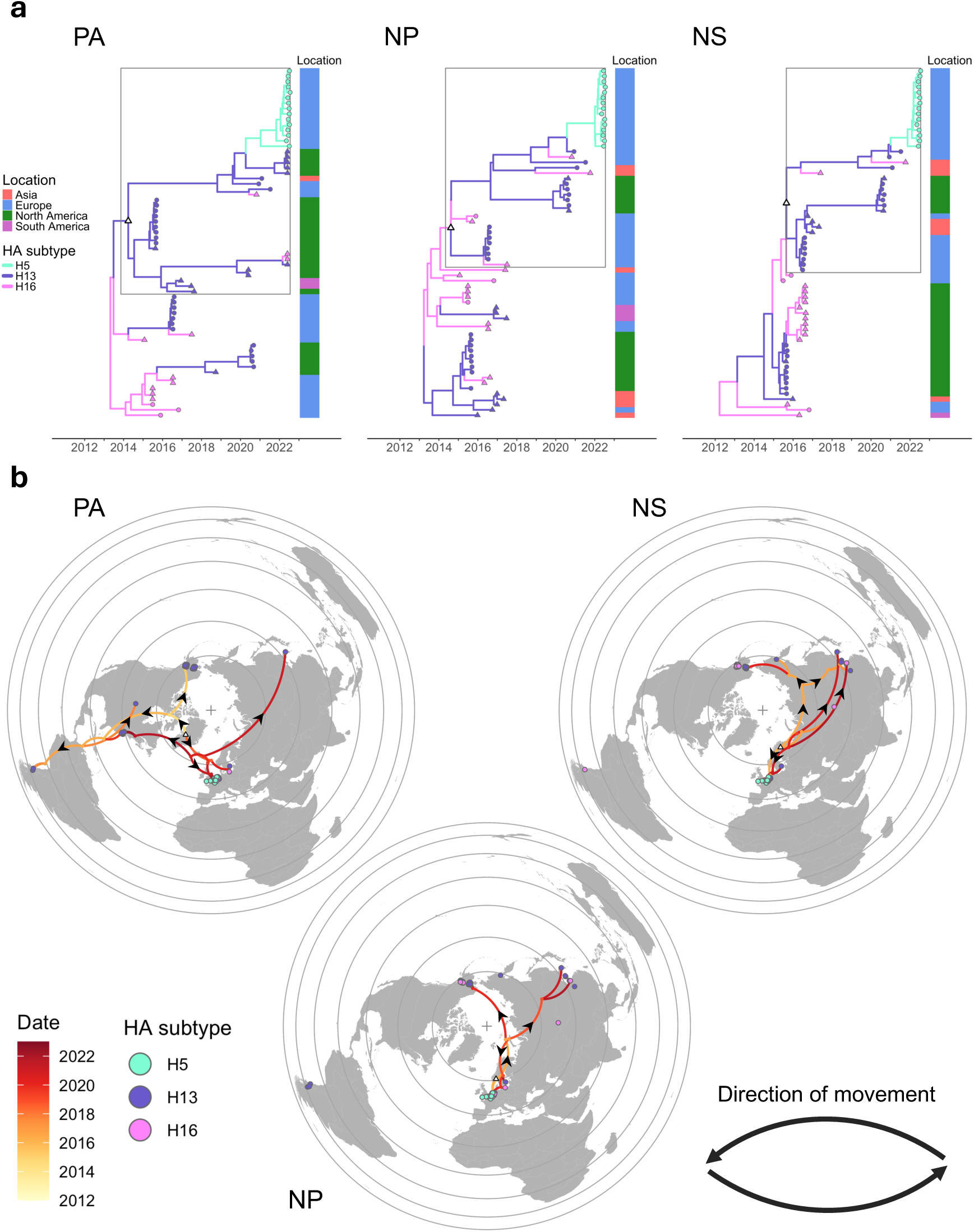
Origins of gene segments acquired by Charadriiformes-adapted H5N1. **a**, Time-scaled PA, NP and NS maximum clade credibility phylogenies consisting of 15 early H5N1/2022/R10 sequences and 50 closely related sequences identified independently for each segment. Branches are coloured by HA subtype and tip shape indicates viruses are present in all three segment phylogenies (circles) or not (triangles). Alongside, a column indicates sampling location. Boxes highlight subtrees below nodes marked by white triangles that are plotted in **b**. **b**, Phylogeographical reconstruction for subtrees highlighted in **a**. with branches coloured by date. Circles indicate locations of sampled viruses are coloured by HA subtype. The direction of movement along curves representing branches of the phylogeny is represented in a counterclockwise direction. Viruses outside selected subtrees are also plotted but are not connected by branches.

Phylogeographical analysis was used to estimate the dates and locations of internal nodes of the phylogenies generated for each segment. For the part of each phylogeny most relevant to the emergence of H5N1 (boxed in Fig 6a), the estimated locations of internal nodes and inferred movements among these and linking them to sampling locations are plotted in Fig 6b (maps with reconstructions for the whole trees in Fig 6a and illustrating uncertainty in the phylogeographical reconstruction are shown in Fig S7). Reconstructions for the segments NP and NS are similar with much of the recent evolutionary history of the estimated to have occurred in northern regions of western Eurasia, with occasional movements eastwards across Eurasia reaching eastern China, Korea and Alaska.

For PA, there is greater evidence of recent dispersal across the North Atlantic, for example, with the Alaskan viruses being most closely related to the PA that emerged in H5N1 being different to those most closely linked by NP and NS. There is also evidence for recent dispersal of PA across the Atlantic with a group of genomes sampled from Charadriiformes in Delaware Bay in May 2022 having PA closely related to that which emerged in H5N1. More generally, these reconstructions demonstrate considerable longitudinal spread of H13 and H16 viruses in Charadriiformes around the globe linking northern regions of Eurasia and the Americas with occasional dispersal to the Pacific region of South America. Migrations of Charadriiformes between coastal breeding sites and pelagic feeding areas can link northern regions where there exist overlapping areas of the various north-south flyways, for example between the East Atlantic and Atlantic (Americas) flyways (4), and therefore Charadriiformes present a potentially important route for spread of AIV between western and eastern hemispheres.

## Discussion

In this study, we describe and quantify dynamics of subclade 2.3.4.4b H5 influenza viruses covering the period in which H5N1 emerged from the previously dominant H5N8 viruses and focusing on the role of reassortment involving H5N1 viruses as they have expanded geographically and diverged genetically. Europe and North America remain the most data-rich regions globally and it is striking that in each of these, subclade 2.3.4.4b viruses independently acquired novel combinations of polymerase and NS segments through reassortment with local LPAI viruses. These regions have acted as largely independent evolutionary systems with their own reassortment dynamics, but in both instances, there is evidence for differential fitness of reassortants and diversification of host specificity. The classification of genomes into reassortant genotypes is broadly consistent with systems presented for the classification of genomes detected in, for example, the UK (42) or Europe (16). As such, our interpretation of results and conclusions are generalisable and not restricted by intricacies of our described approach. Predictions of the medium-to-longer term fate of reassorted lineages is complicated by the unknown phenotypic impacts of further future reassortment events. While the panzootic continues in various geographies and host types, reassortment opportunities will continue to generate viral genomic diversity that is both intriguing from a virological perspective and concerning for public and animal health.

We have used a combination of phylodynamic and experimental approaches to explore the fitness of reassortant lineages of 2.3.4.4b HPAI viruses in different hosts. These approaches are complementary and together can provide a richer understanding of complex dynamics. For example, the lack of NS1-mediated host shutoff mediated by the H5N1/2022/R10 (EA-2022-BB) representative in cells from water- and land-fowl could explain the apparent low fitness of this virus in these hosts as inferred from phylodynamic analyses. Seemingly incongruous results may also emerge with, for example, replication kinetics in cell lines and polymerase assays suggesting H5N1/2020/R1 (EA-2020-C) to be superior to H5N1/2021/R2 (EA-2021-AB), however LBI indicate that the latter emerged with a transmission advantage before replacing the former in Europe. It is possible that H5N1/2021/R2 surpassed H5N1/2020/R1 in a trait not measured experimentally, such as the persistence of the virus in the environment, or perhaps H5N1/2020/R1 exhibited increased replication leading to elevated virulence resulting in a transmission cost. There are limitations in that our assays do not measure all aspects of phenotype that contribute to viral fitness while metrics like LBI and phylogenetic persistence are affected by biases in sampling intensity in different geographies and host types.

Further there are likely gaps in analysis through unsampled ancestry across the different genotypes. Indeed, due to the collection and sampling methods for dead wild birds there is likely to be extensive unsampled ancestry during outbreaks. Phylodynamic metrics aim to account for this by incorporating genetic diversity and so are far more robust than case numbers alone. However, there may also be biases in terms of which infected birds are sampled, perhaps towards sick or dead birds or those that live in groups where die-offs are more apparent or easier to sample. As such, the role of important hosts that may be under-sampled, perhaps being less likely to exhibit severe disease or to be discovered, could be underestimated. In related work, we are using both infection and bird population distribution data and machine learning approaches to address this issue (43).

Exploring evolutionary dynamics in Europe in greater detail, we highlight the emergence of the H5N1/2022/R10 lineage through reassortment around April 2022 with five segments inherited from an H5N1/2021/R2 (EA-2021-AB) virus and the remaining three likely from an H13 LPAI virus circulating in Charadriiformes. After emergence, this reassortant lineage appears to have circulated in a somewhat exclusively within Charadriiformes, a distinct host community with its own evolutionary dynamics and seasonality. We show that host-dependent LBI and phylogenetic persistence estimates indicate this lineage to be specialised for transmission in Charadriiformes with lower fitness and transmission potential in other avian hosts. In laboratory studies, we have demonstrated that this lineage exhibits similar replication kinetics and polymerase function compared with the H5N1/2021/R2 representative. However, we also show this lineage to lack effective NS1-mediated host translational shutoff in cells of Galliformes or Anseriformes origin. If this corresponds to an inability of H5N1/2022/R10 viruses to modulate host gene expression in birds of these avian orders, this could contribute to their lower epidemiological fitness in these hosts, relative to the host-generalist H5N1 viruses from which they descended. It was not possible for us to test these viruses using cell lines originating from Charadriiformes due to a lack of availability, though this would be an interesting area for further work. More generally, it would be interesting to elucidate why H13 and H16 viruses are endemic to Charadriiformes (1,3,41).

During the ongoing H5N1 panzootic, there is further evidence for host diversification with at least one other instance in which H5N1 viruses have begun to transmit in a novel host population with very limited overlap with the typical community of hosts, the ongoing epizootic in dairy cattle within the USA. Diversification of host specificity has various interesting evolutionary potential consequences. When viral transmission in different host types becomes increasingly separated, competition between lineages is diminished according to the extent to which lineages cease to transmit within common populations. With lineages no longer in direct competition for hosts, they can evolve independently in parallel, likely with adaptive changes reflecting the different host associations, increasing the overall genomic diversity of subclade 2.3.4.4b viruses. Through independent evolution in different host types, they are exposed to distinct selective pressures and novel reassortment opportunities. Viruses are expected to gain adaptive mutations and may acquire genomic segments from influenza viruses that are endemic to different host types, both of which have the potential to further accentuate differences in host specificity. It is vital to understand how the evolution of these virus lineages in different fitness landscapes could affect their potential to transmit to humans or other mammals.

There is currently significant attention being given to identifying the drivers and barriers to geographical spread of subclade 2.3.4.4b HPAI viruses, with a view to predicting the areas in which the risk of wild bird infections is elevated and evaluating the risk of transmission at the wildlife-livestock interface. In addition to such general objectives, stakeholders and policy makers seek answers to specific questions, for example, concerning the probability that the 2.3.4.4b lineage circulating in US dairy cattle could be spread to Eurasia by wild birds. At present, the geographical spread of this lineage remains restricted to the USA and there is no evidence of sustained transmission of 2.3.4.4b HPAI following dispersal from North America to Eurasia, though we show North America and Eurasia to be well connected through phylogeographical reconstructions of H13 and H16 LPAI associated with Charadriiformes. Modelling efforts in this field that use, for example, data on the distributions and migratory patterns of wild birds or indeed other concerned animals should be aware of differences in host specificity between lineages and incorporate information on the host-specific transmission potential of different lineages where possible.

## Materials and Methods

### Data

All influenza A H5 genomes with sequence data for all eight segments and submission dates between 1^st^ January 2020 and 9^th^ November 2024 were downloaded from GISAID (39). Genomes resulting from environmental sampling or inferred to be laboratory-derived reassortant viruses were excluded, or with incomplete information for collection date. Furthermore, genomes from avian hosts that lacked sufficient information to derive avian order, for example those with imprecise host information such as ‘avian’ or ‘wild bird’ were excluded. Gene sequences with high nucleotide similarity to the PA, NP and NS segments of the virus A/gull/France/22p015977/2022 (GISAID accession number: EPI_ISL_13519451) were identified using BLAST (34), performed on 4^st^ October 2023 on the GISAID platform (39).

### Alignment and clustering

For each gene segment, nucleotide sequences were aligned using MAFFT v7.511 (44). Alignments were trimmed to coding regions and insertions of a non-multiple of three bases or insertions present in fewer than 10% of sequences were deleted. Genomes with nucleotide coverage of less than 85% in any of the eight gene segments were excluded from further analysis. For each segment, a set of unique nucleotide sequences was identified, and every isolate was associated with a representative sequence in this set. From each set of unique sequences, a phylogeny was constructed using a general time reversible (GTR) model of nucleotide substitution using FastTree v2.2.0 (45,46). The resulting phylogenies were midpoint rooted and for each segment, clusters were defined upon the relevant tree. Within each bifurcating tree, each node, *i*, was considered in relation to its parental node, *i*_*p*_, and its sibling node, *i*_*s*_, with respect to a threshold branch length, *τ*. A novel cluster label was assigned to node *i* dependent on the branch length of the node, *b*_*i*_, and the isolation of node *i*, *I*_*i*_, defined as the sum of branch lengths *b*_*i*_, 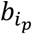 and 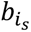. A novel cluster was assigned when *b*_*i*_ < *τ*’ and *I*_*i*_ < 3 *τ*’ where *τ*’ is *τ* adjusted for depth in the phylogeny according to

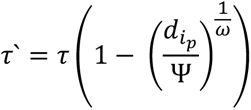

where the depth of the parental node, 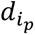, in the phylogeny was defined as the smaller sum of branch lengths within the two subtrees descended through nodes *i* and *i*_*s*_, and where Ψ is the total sum of branch lengths in the tree. When implemented with *ω* = 1, a mild tendency to define new clusters deeper in the phylogeny is encouraged, whereas higher values would increase this tendency. This approach was implemented with a threshold branch length, *τ*, of 0.0075 substitutions per site for segments PB2, PB1, PA, HA and NP and 0.001 for the shorter segments NA, M and NS, using code available at https://github.com/will-harvey/toolkit_seqTree. This process resulted in a numeric cluster ID for each gene segment where 1 was assigned to the most frequently detected cluster and increasing values represented increasingly less populated clusters. For each genome, the combination of cluster IDs for each of the eight gene segments provided a profile of cluster IDs in order of decreasing segment size: PB2_PB1_PA_HA_NP_NA_M_NS. For each profile with at least ten assigned genomes, a cluster label was also generated. Each label was generated to indicate virus subtype, the year in which a genome assigned to profile was first sampled, and an emergence, or ‘R’, number which refers to the position of the profile in the chronological sequence, within that year, in which novel profiles were first sampled and made publicly available according to the earliest collection date on the GISAID platform. To enable comparison with earlier iterations of this approach, cluster IDs were also generated per-segment directly from aligned nucleotide sequences using a hierarchical clustering of pairwise nucleotide distances implemented with a nearest neighbour algorithm using the R package *bioseq* (47,48). Corresponding genotype labels from the EURL Genein2 system (https://github.com/izsvenezie-virology/genin2) were identified. Corresponding genotype labels from the USDA GenoFLU system (20) were accessed directly from GISAID. Here, genotypes were reported as corresponding in Table 1 if that genotype represented more than 5% of sequences assigned to a reassortant type in this work.

### Phylogenetic analysis

Time-resolved phylogenetic trees were generated from nucleotide sequences with associated sampling dates using BEAST v1.10.4 (49). Sequence evolution was modelled using a Shapiro-Rambaut-Drummond-2006 (SRD06) model (50), an extension of the Hasegawa-Kishino-Yano substitution model (51) with a gamma distribution with four categories describing among-site variation (52) (HKY + Γ_4_) that also allows substitution rate parameters and the rate heterogeneity model to vary at the third codon position relative to the first two. This model was implemented with a relaxed molecular clock (53) with branch rates drawn from a lognormal distribution and a Bayesian skygrid coalescent model (54) which eschews any prior assumption on changes in effective population size through time. Convergence, effective sample sizes, and mixing properties were inspected using Tracer v.1.7.2 (https://github.com/beast-dev/tracer/releases). Independent chains were combined using LogCombiner v1.10.4 (49) and maximum clade credibility (MCC) trees were identified using TreeAnnotator v1.10.4 (49). For the large phylogeny in Fig 2, a posterior sample of trees was generated using Delphy v1.0.3 (https://github.com/fathominfo/delphy-web) with sequence evolution modelled using a GTR model with site heterogeneity drawn from a continuous gamma distribution. Delphy was run until the minimum effective sample size exceeded 200 and the MCC tree was identified using TreeAnnotator. Phylogenies were visualised in *R* (47) using the *ggtree* package (55).

### Discrete trait analysis

Rates of transition between geographical regions, host types and subtypes were estimated in BEAST v1.10.4 (49) using asymmetric discrete-state continuous Markov chain (CTMC) models of trait evolution that allowed rates of transition between any pair of traits to differ depending on the direction (56). Bayesian stochastic search variable selection was used to identify phylogenetically robust transitions. This involves the inclusion of a binary indicator variable associated with each transition. This indicator variable was sampled by MCMC and transitions that were included in the model (indicator variable = 1) in more than 50% of MCMC samples were considered phylogenetically robust. For such transitions, conditional mean rates (i.e. posterior mean averaged across samples where indicator variable = 1) were calculated.

### Continuous phylogeography

Spatial diffusion was mapped to viral phylogenies using the continuous phylogeographical framework implemented in BEAST (57). Geographical coordinates were estimated from publicly available location. To facilitate the visualisation of diffusion in regions surrounding the around the North Pole, coordinates were transformed to the Lambert azimuthal equal-area projection centred on the North Pole using the *mapproj* package (58) in *R* (47). Spatial diffusion was modelled with a relaxed random walk model with branch-specific scaling factors drawn from a Cauchy distribution which has been shown to be well suited to capturing heterogeneity in rate diffusion for avian influenza (59). This was implemented with a distribution of 901 empirical trees sampled from the posterior of a BEAST analysis run without traits, with an HKY + Γ_4_ substitution model, relaxed molecular clock model with rates drawn from a lognormal distribution and a Bayesian skygrid coalescent model.

### Local branching index

The local branching index (LBI) (60) was calculated for both time-scaled and non-scaled phylogenies. To monitor LBI through time, LBI was repeatedly recalculated for a series of temporal slices of lineages. For each time window, a subtree consisting of only ancestral lineages was identified, a procedure which alters the branching pattern such that only tips within a particular time window exist. LBI was then calculated using the branching pattern within this subtree. To calculate host-specific LBI, subtrees consisting of branches that descend tips of a given host type were identified and the metric was recalculated using the revised branching structure. The local branching index implemented in R (47) using code available at https://github.com/will-harvey/toolkit_seqTree.

### Multi-type birth death model

The multitype-tree birth-death model (61) implemented in BEAST2 v2.6.3 (62) was used to estimate the effective reproductive number (*R_e_*). This was used to estimate whether genotypes, defined prior to the analysis, exhibited variation in *R_e_* through time (63). The following priors were specified: 1) *R_e_*: a mean of *R_0_* 2.5 with 95% HPD (0.6, 6), and were estimated over five equidistant time intervals depending on the size of the overall tree; 2) the “becomeUninfectiousRate”, which refers to the number of days from infection to recover: a mean of 52 (365/7 days) with 95% highest density interval [4.44, 224] (equivalent to [1.63, 82.21] infectious days) under a lognormal distribution.; 3) the sampling portion, a mean of 2×10^−4^ with 95% HPD (1×10^− 5^, 1×10^−3^); 4) the substitution rate: a mean of 0.003 and standard deviation of 0.001 under a lognormal distribution and 5) the origin time of the epidemic: the estimated time to the most recent common ancestors (tMRCAs) of the subsampled HA gene tree with priors described above.

### Cells and viruses

Madin-Darby canine kidney cells (MDCK) (American Type Culture Collection, ATCC), human embryonic kidney cells (293T) (ATCC), pekin duck embryo fibroblasts (CCL-141, kindly provided by Dr Leah Goulding) and Japanese quail Fibrosarcoma Cells (QT-35, kindly provided by Dr Laurence Tiley) were cultured in Dulbecco’s modified Eagle’s medium (DMEM, Sigma) supplemented with 10% foetal bovine serum (FBS, Gibco), 2 mM glutamine, 100 U/mL penicillin and 100 µg/mL streptomycin. Chicken embryo fibroblasts (DF1) were cultured in DMEM:Nutrient Mixture F12 (DMEM F12, Sigma) supplemented with 10% FBS, 2% chicken serum, 2 mM glutamine, 100 U/mL penicillin and 100 µg/mL streptomycin. Chicken lung epithelial cell (CLEC213 (64), kindly gifted by Sascha Trapp) were cultured in DMEM F12 supplemented with 8% FBS, 2 mM glutamine, 100 U/mL penicillin and 100 µg/mL streptomycin. Human-engineered haploid cells (eHAP, Horizon Discovery) with ANP32A, ANP32B and ANP32E genes knock-out (tKO) by CRISPR-Cas9 as described previously (65), were cultured in Iscove’s modified Dulbecco’s media (IMDM, Gibco) supplemented with 10% FBS, 100 U/mL penicillin, 100 µg/mL streptomycin and 1x Non-essential amino acids (NEAA, Gibco). All cell lines were cultured at 37-39°C with 5% CO_2_, regularly tested for mycoplasma contamination and passaged twice weekly.

All reverse genetics work was carried out under a license (GMRA1811) from the UK Health & Safety Executive. A/chicken/England/053052/2021 (H5N1) (AIV07), A/chicken/Scotland/054477/2021 (H5N1) (AIV09), and A/chicken/England/085598/2022 (H5N1) (AIV48) harbouring HA and NA genes of the laboratory adapted strain A/Puerto Rico/8/1934 H1N1 (PR8) were generated by reverse genetics as previously described (66). Briefly, 2 × 10^6^ 293T cells were transfected with pHW2000 reverse genetics plasmids (250 ng of plasmid for each virus segment) in OptiMEM using 4 µL of Lipofectamine2000 according to the manufacturer’s instructions. Twenty-four hours post-transfection, media was replaced with serum-free DMEM supplemented with 0.14% fraction V bovine serum albumin (BSA) (w/v) and 1 μg/mL of L-(tosylamido-2-phenyl) ethyl chloromethyl ketone (TPCK)-treated trypsin. Virus-containing supernatant was collected after a further 2-day incubation and clarified by centrifugation. 10 µL of clarified supernatants were used to infect 10-day-old embryonated hen’s eggs which were incubated for 2 days at 37°C. Following a further 4°C overnight humane killing, allantoic fluids were collected, clarified and stored until titrated by plaque assay and further usage.

### Virus quantification by plaque assay

Confluent monolayers of MDCK cells overexpressing galline ANP32A (kindly provided by Prof Massimo Palmarini) (1 × 10^6^ seeded the day before infection in 12-well plates) were washed once with PBS and infected with tenfold serial dilutions of virus. Following a 1 h incubation at 37 °C, cells were overlaid with DMEM supplemented with 0.14% fraction V BSA, 1 µg/mL TPCK-treated trypsin and 1.2% Avicel and incubated for 2 days at 37 °C, 5% CO_2_. Cells were fixed with 10% neutral buffered formalin and stained with a 1% Toluidine Blue solution for at least 1 h. Staining solution was rinsed under tap water, plates were air-dried and plaques were counted.

### Growth kinetics analysis

Monolayers of CLEC213, DF1, CCL-141 and QT35 cells were washed once with PBS and infected at multiplicity of infection (MOI) 0.001 with virus diluted in serum-free medium for 1 h at 37 °C. Inoculum was replaced with serum-free medium supplemented with 0.14% BSA and 1 µg/mL TPCK-treated bovine pancreas trypsin. Supernatants were collected at various times post-infection and stored at −80°C until infectious titres were determined by plaque assay.

### Measurement of type I IFN induction

Sub-confluent monolayers of CLEC213 were co-transfected with 500ng of segment 8-encoding pHW2000 plasmids and 100 ng of reporter plasmid using 1μL of Lipofectamine2000. The following day, cells were transfected with 1 µg of Poly I:C using 2 μL of Lipofectamine2000 and lysed with 1x Reporter lysis buffer a further 24h post-transfection. Luminescence levels were measured using 40 µL of lysate and 25µL of luciferase assay reagent (Promega) in a Biotek citation 3 imaging reader using the Biotek Gen5 software.

### Measurement of host protein synthesis shutoff

Sub-confluent monolayers of CLEC213, DF1, CCL-141 and QT35 cells were co-transfected with 500 ng of segment 8-encoding pHW2000 plasmids and 100 ng of pRL reporter plasmid using 1 μL of Lipofectamine2000. Luminescence was measured using a *Renilla* Luciferase Assay kit 48h post-transfection. Briefly, cells were lysed with 100μL of 1× passive lysis buffer and frozen overnight at −20°C. Cells were scraped off the plate, and lysates were clarified by centrifugation (8000rpm, 5 minutes, 4°C), 40 μL of each clarified supernatant was transferred to a white-bottomed 96-well plate and luminescence was measured in a Biotek citation 3 imaging reader, injecting 25 μL of 1× *Renilla* luciferase substrate and using the Biotek Gen5 software.

### Polymerase activity assays

eHAP tKO cells were transfected in 24-well plates with Lipofectamine 3000 transfection reagent (Thermo Fisher) with the following amounts of pCAGGs expression plasmids: 160 ng PB2, 160 ng PB1, 160 ng PA, 160 ng NP, 160 ng *Renilla* luciferase, 320 ng species-specific polI vRNA Firefly luciferase and 160 ng ANP32A or empty pCAGGs. CLEC213, DF1 and CCL-141 cells were transfected in 12-well plates with double the amount of plasmids in the same ratios as above. However, these cells were not transfected with ANP32A plasmids. eHAP tKOs were transfected with human pHOM1-Firefly plasmid and avian cells were transfected with pCOM1-Firefly, as previously described (66). Cells were lysed 24h post-transfection using passive lysis buffer (Promega) and polymerase activity measured using the Dual-luciferase Reporter assay system (Promega) and a VANTAstar plate reader (BMG labtech). Firefly signal was normalised to *Renilla* signal to give relative luminescence units (RLU).

### Immunoblotting

Cells were lysed in radioimmunoprecipitation (RIPA) buffer supplemented with an EDTA-free protease inhibitor or Leammli’s buffer, heated at 95°C for 5 minutes and subjected to polyacrylamide gel electrophoresis and transferred to polyvinylidene difluoride membranes. Membranes were blocked in PBS/0.2% Tween20(TBST)/5% milk for 1h at room temperature and incubated overnight at 4°C with the following primary antibodies: mouse FLAG (Sigma, F1804, 1:250), rabbit PB2 (GeneTex, GTX125926, 1:500), PA (GeneTex, GTX118991, 1:500), mouse α-Tubulin (Abcam, ab7291, 1:1250), rat α-Tubulin (LifeTechnologies, MA180017, 1:2000), rabbit NS1 (V29, homemade, 1:500). Following three 5-minute TBST washes, membranes were incubated with the following near-infrared fluorescent secondary antibodies: goat anti-mouse IgG IRDye 800CW (Abcam, ab216772), goat anti-rabbit IgG IRDye 680RD (Abcam, ab216777, 1:10,000), goat anti-rat IgG IRDye 680RD (LI-COR, 926-68076, 1:10,000) and goat anti-rabbit IRDye 800CW (LI-COR, 926-32211, 1:10,000) for 1h at room temperature. Following three PBST washes, membranes were imaged using an Odyssey DLx or an Odyssey Fc imaging systems (Li-Cor Biosciences) using the Image Studio Lite software.

## Acknowledgements

We gratefully acknowledge all data contributors, i.e., the Authors and their Originating laboratories responsible for obtaining the specimens, and their Submitting laboratories for generating the genetic sequence and metadata and sharing via the GISAID Initiative, on which this research is based.

## Funding Statement

We acknowledge support for this research consortium from the Medical Research Council (MRC, UK), Biotechnology and Biological Sciences Research Council (BBSRC, UK) and Department for Environment, Food and Rural Affairs (Defra, UK) as ‘FluMAP’ (grant number BB/X006204/1, BB/X006166/1), ‘FluTrailMap’ (BB/Y007271/1, BB/Y007298/1) and FluTrailMap-One Health (MR/Y03368X/1). Furthermore, this research was funded by the BBSRC via Institute Strategic Grants to the Roslin Institute (BBS/E/RL/230002C and BBS/E/RL/230002D) and the Pirbright Institute (BBS/E/PI/230002A and BBS/E/PI/230002B), via Evolution and Ecology of Infectious Diseases grant (BB/V011286/1), and through NIFA-USDA (USA), by the European Union’s Horizon 2020 research and innovation programme (Grant no. 874735), and by Defra (UK) as part of the International Coordination of Research on Infectious Animal Diseases (ICRAD) consortium (contract no. SE2223, FluSWITCH), and by project numbers SE2230 (iPrepare), SV3400, SV3032, and SV3006. The funders had no role in study design, data collection and analysis, decision to publish, or preparation of the manuscript.

## Supplementary Information

**Fig. S1.**
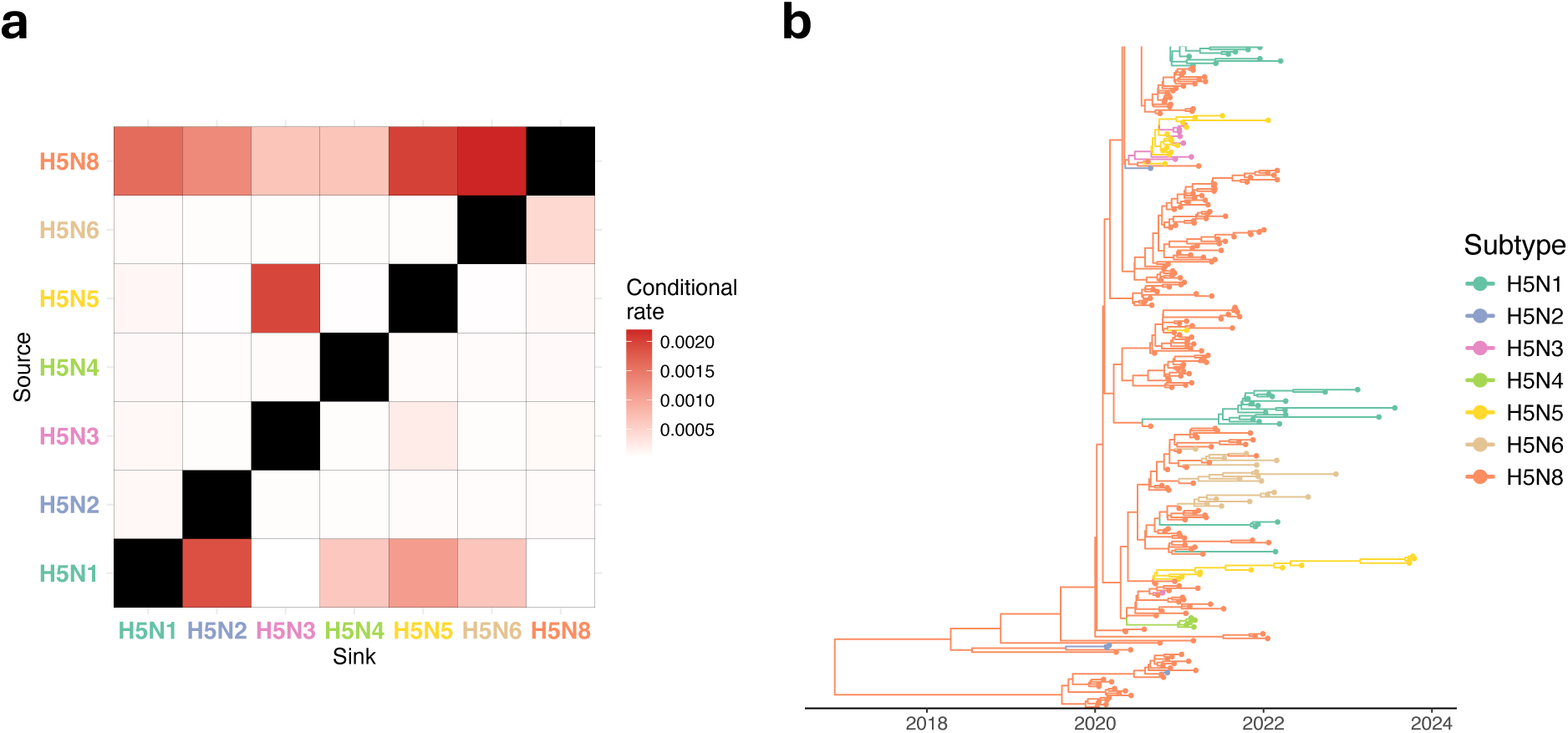
Transitions between subtypes in phylogeny displayed in Fig 1. **a**, Rates of transition between subtypes estimated using asymmetric discrete-state continuous Markov chain (CTMC) model with Bayesian stochastic search variable selection (BSSVS) was used to identify phylogenetically robust transitions. Mean rates conditional on BSSVS indicator rates are shown. **b**, Close-up focusing on the part of phylogeny in Fig 1 in which several reassortment events resulting in changes from H5N8 to other subtypes are inferred.

**Fig. S2.**
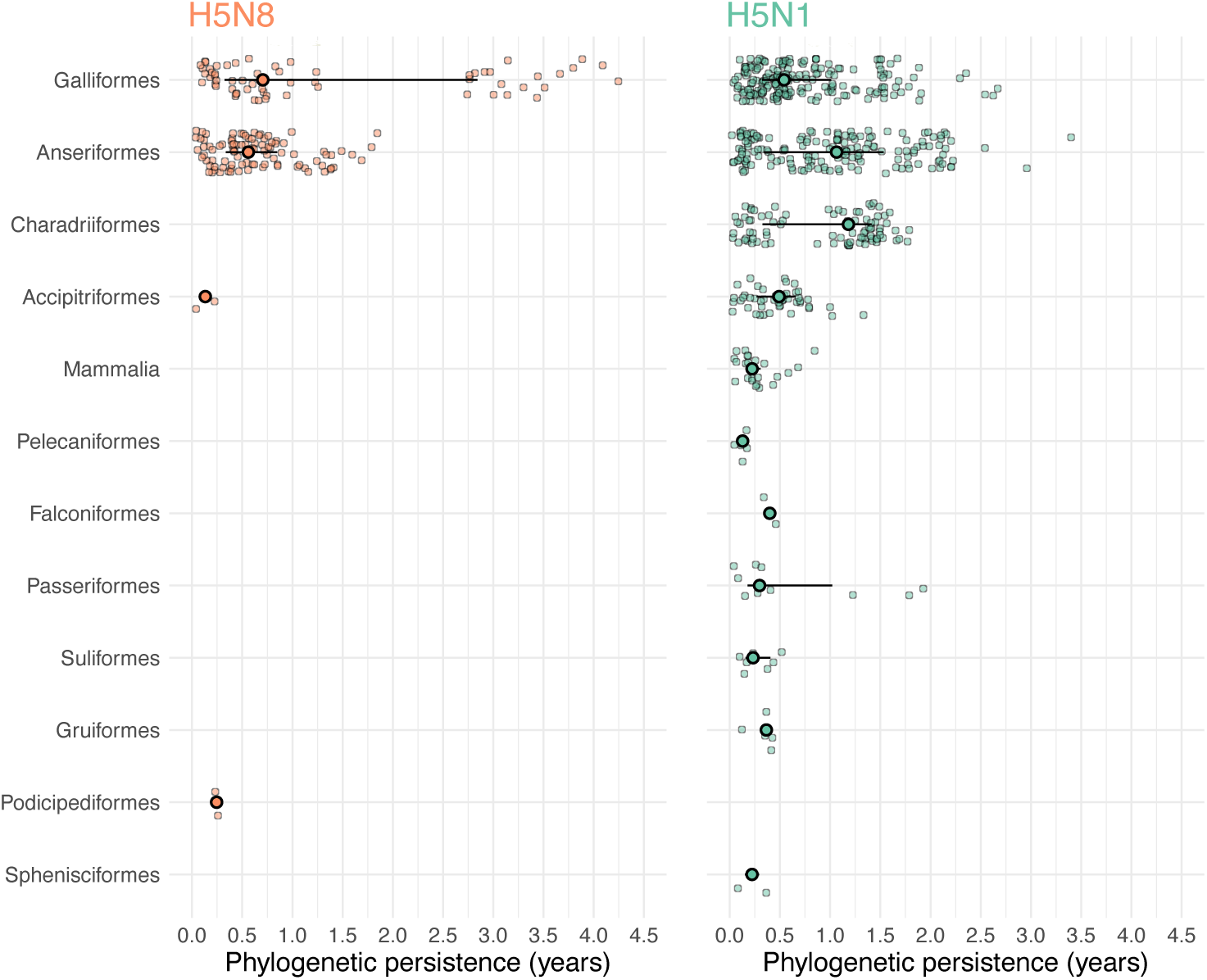
Phylogenetic persistence in different host types by subtype, covering the period of emergence of panzootic H5N1. Duration of host-specific phylogenetic persistence for avian orders and for mammals. Small circles indicate individual measurements from across the phylogeny and to aide visualisation of overlapping points, random adjustment in the vertical dimension has been added. The positions of large, opaque circles indicate median persistence with lines showing inter-quartile range.

**Fig. S3.**
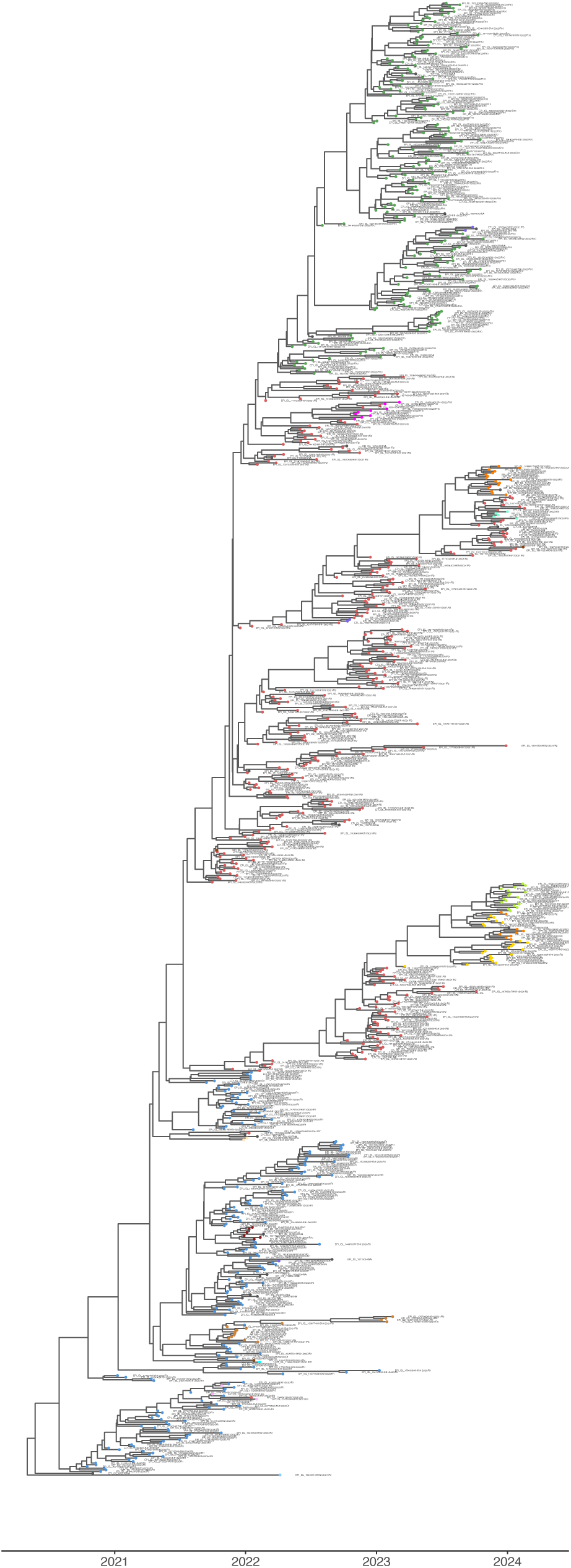
Fully annotated version of time-scaled HA phylogeny shown in Figure 3a featuring sample of viruses detected in Europe. Tip points are coloured by reassortant genotype and tree can be zoomed to read tip labels that denote GISAID isolate ID and genotype label.

**Fig. S4.**
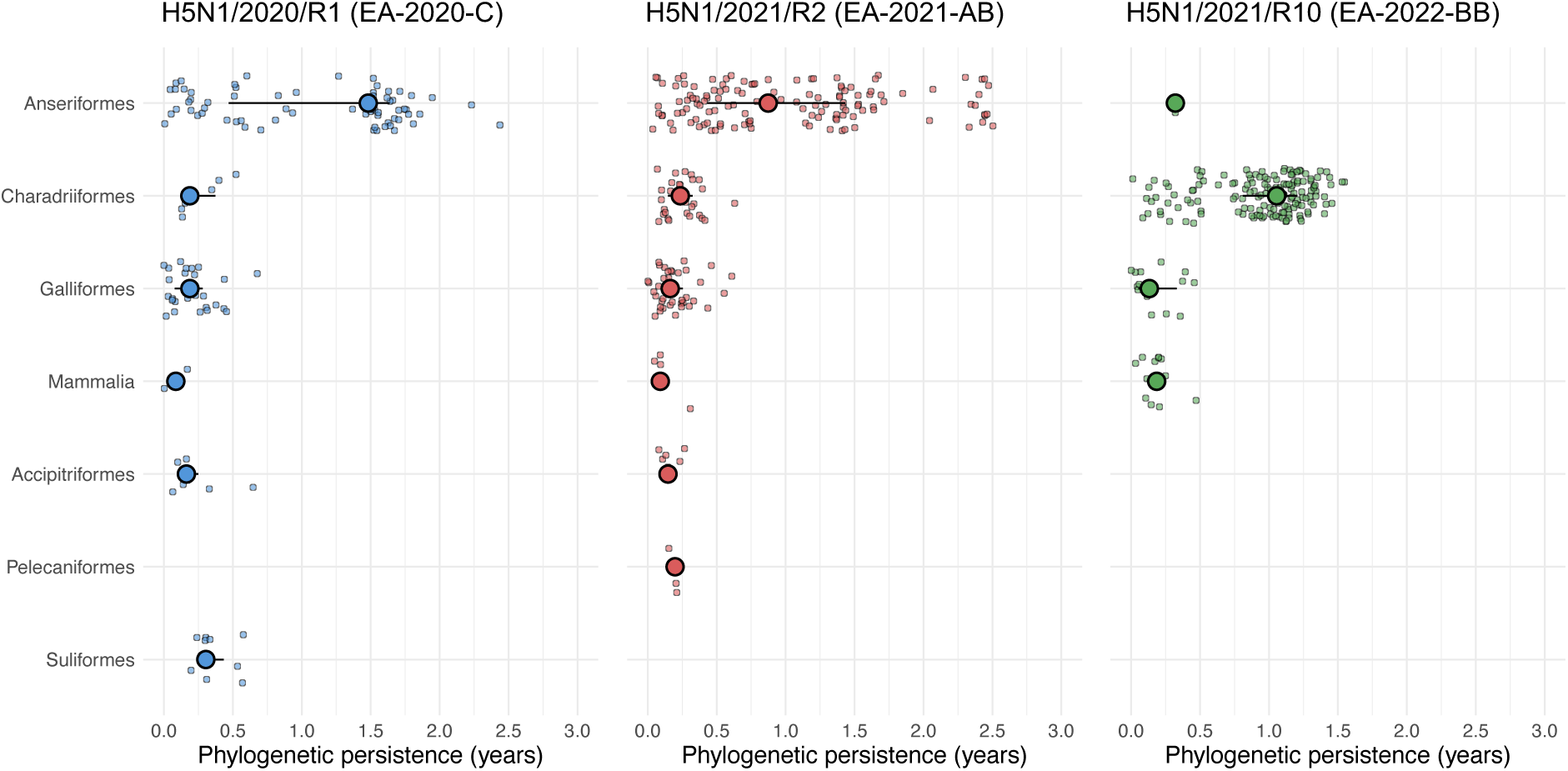
Phylogenetic persistence in different host types by H5N1 genotype, covering the period October 2020 to February 2024 in Europe. Duration of host-specific phylogenetic persistence for avian orders and for mammals. Small circles indicate individual measurements from across the phylogeny and to aide visualisation of overlapping points, random adjustment in the vertical dimension has been added. The positions of large, opaque circles indicate median persistence with lines showing inter-quartile range.

**Fig. S5.**
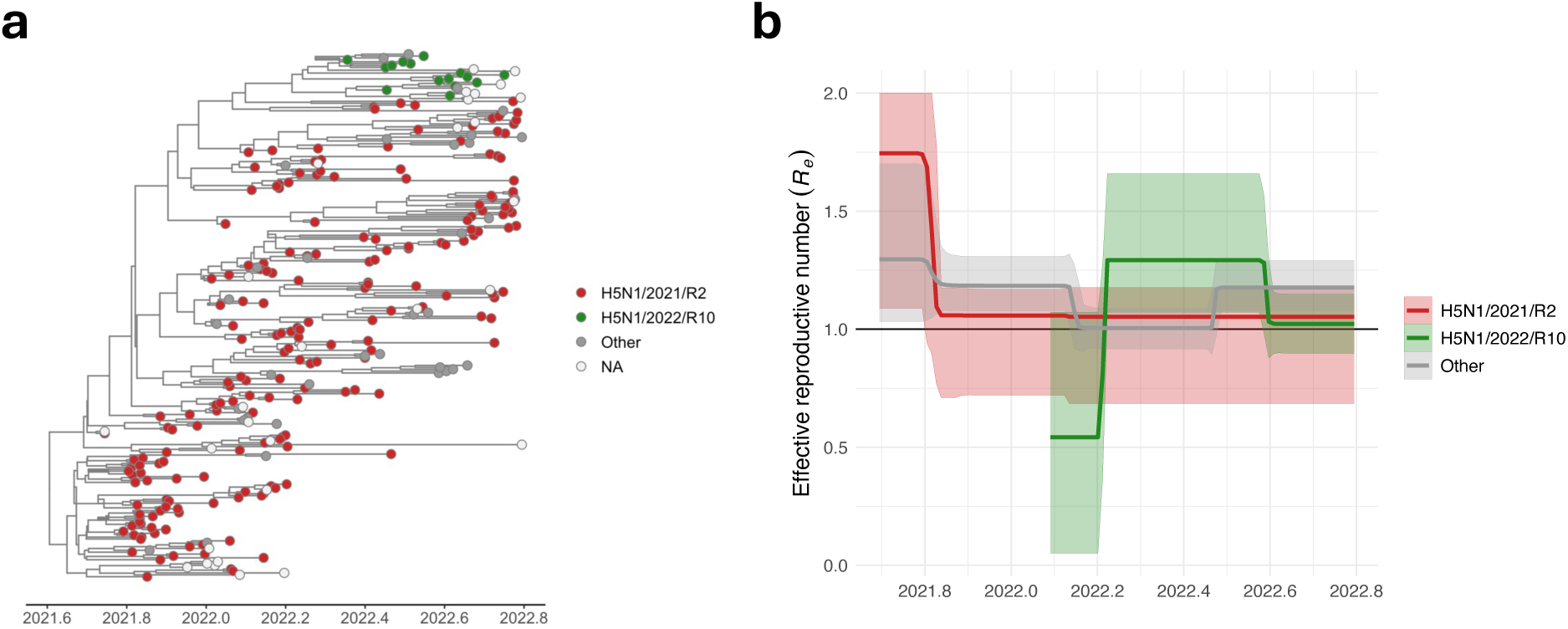
Multi-type birth death model analysis covering the emergence of H5N1/2022/R10 from H5N1/2021/R2. **a**, Time-scaled HA phylogeny showing stratified sample of clade 2.3.4.4b H5N1 viruses sampled between 30^th^ September 2021 and 18^th^ October 2022. **b**, Effective reproductive number (*R_e_*) through time estimated for predefined groups in phylogenetic analysis.

**Fig. S6.**
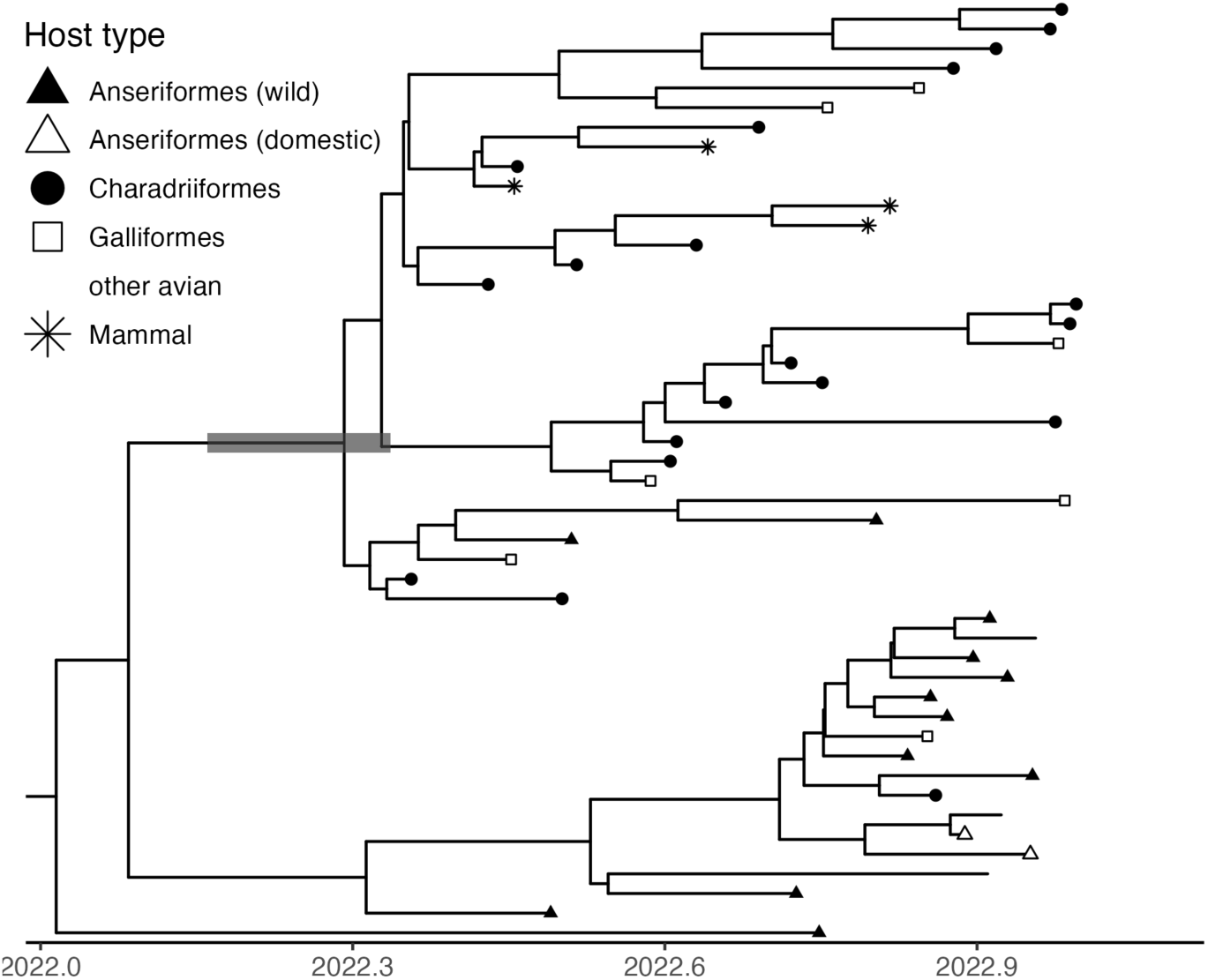
Emergence of genotype H5N1/2022/R10. Time-scaled HA phylogeny showing stratified sample of early H5N1/2022/R10 viruses sampled in 2022 and closely related H5N1/2021/R2 viruses with host type indicated by tip shape according to the key. A grey bar indicates the 95% HPD estimate for the date of tMRCA of the emergent H5N1/2022/R10 clade, 3 April 2022 (95% HPD, 28 Feb-3 May).

**Fig. S7.**
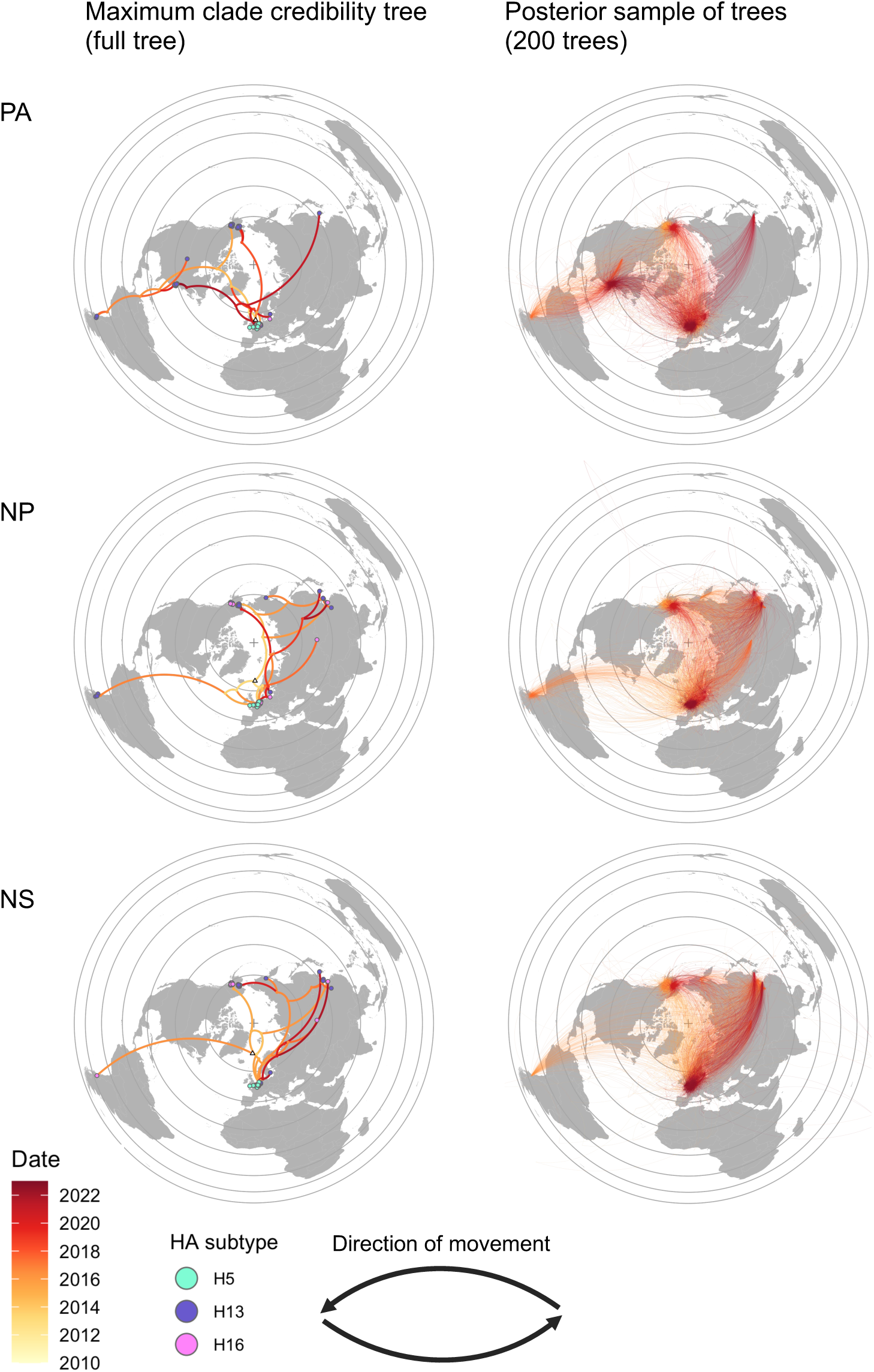
Phylogeographical reconstructions of gene segments acquired by H5N1/2022/R10. **a**, Phylogeographic reconstruction for full time-scaled PA, NP and NS maximum clade credibility phylogenies show in Fig 1a. Each phylogeny consists of 15 early H5N1/2022/R10 sequences and 50 closely related sequences identified independently for each segment. Branches are coloured by date, coloured circles indicate locations of sampled viruses are coloured by HA subtype, and a white triangle shows the estimated location of the root. The direction of movement along curves representing branches of the phylogeny is represented in a counterclockwise direction **b**, Superimposition of 200 phylogenetic trees randomly sampled from the posterior illustrating uncertainty in the phylogeographical analyses.

**Table S1.**
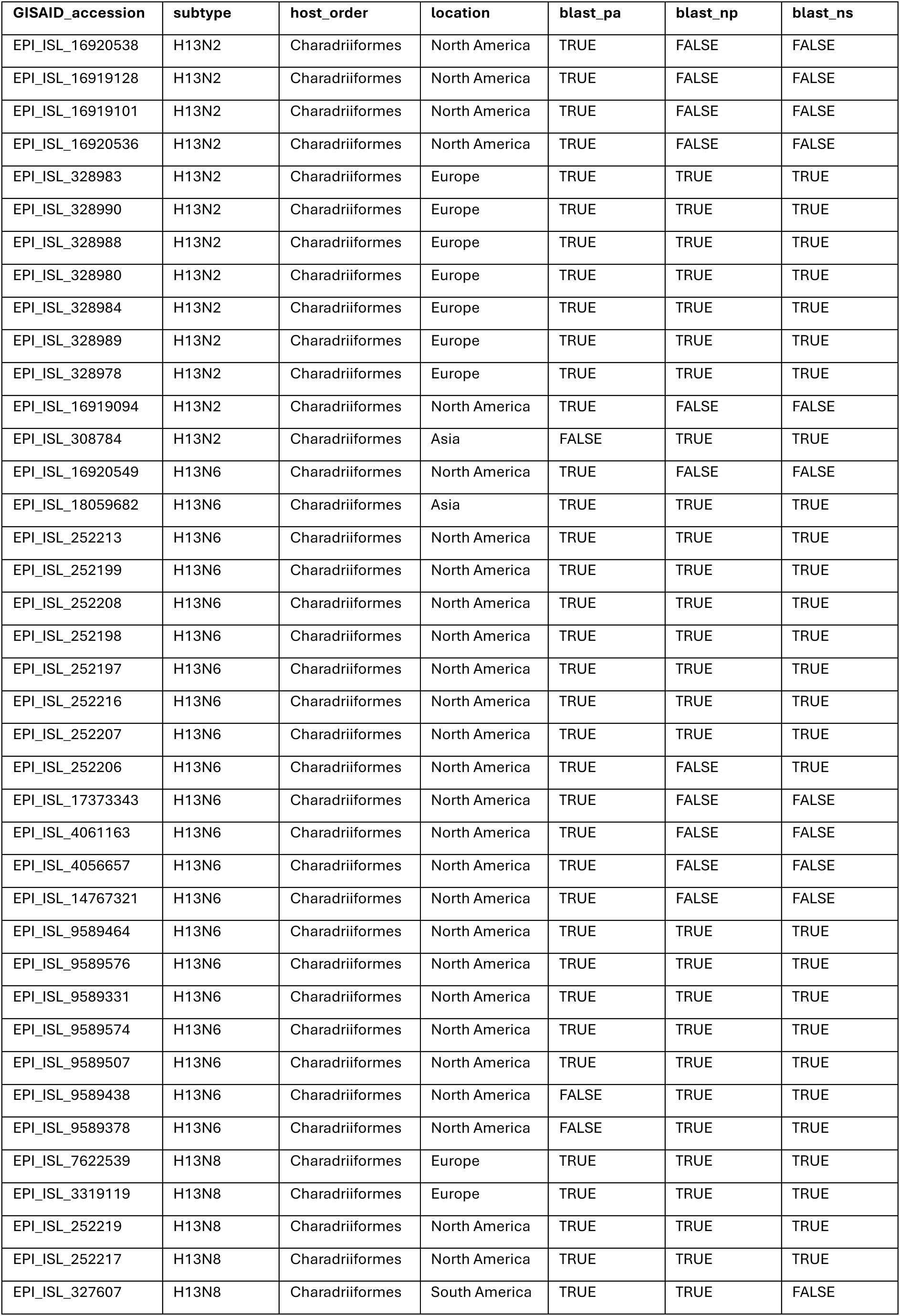
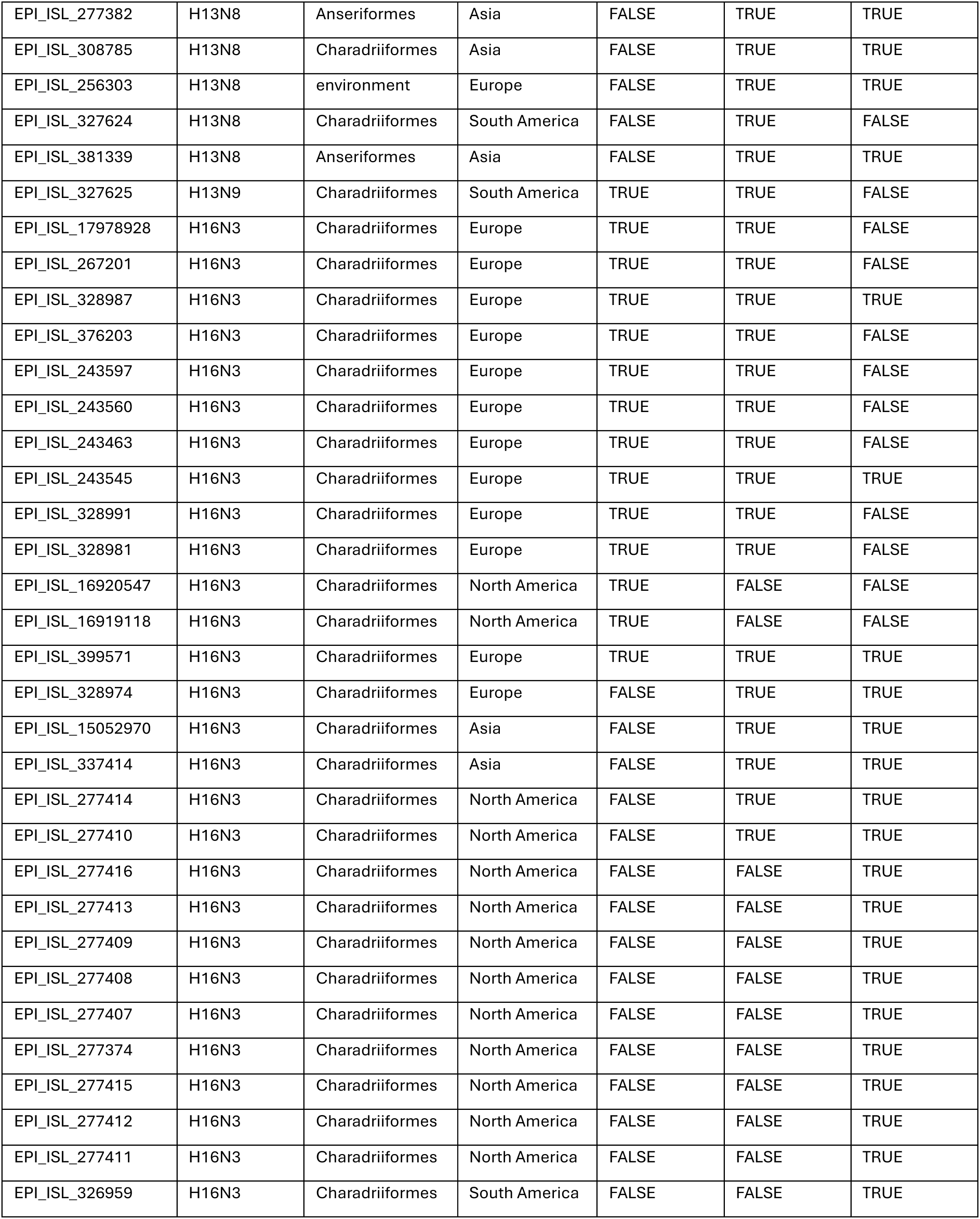
GISAID isolates identified in BLAST searches based on the PA, NP and NS segments of virus A/gull/France/22p015977/2022 (GISAID accession: EPI_ISL_13519451).

| GISAID_accession | subtype | host_order | location | blast_pa | blast_np | blast_ns |
| --- | --- | --- | --- | --- | --- | --- |
| EPI_ISL_16920538 | H13N2 | Charadriiformes | North America | TRUE | FALSE | FALSE |
| EPI_ISL_16919128 | H13N2 | Charadriiformes | North America | TRUE | FALSE | FALSE |
| EPI_ISL_16919101 | H13N2 | Charadriiformes | North America | TRUE | FALSE | FALSE |
| EPI_ISL_16920536 | H13N2 | Charadriiformes | North America | TRUE | FALSE | FALSE |
| EPI_ISL_328983 | H13N2 | Charadriiformes | Europe | TRUE | TRUE | TRUE |
| EPI_ISL_328990 | H13N2 | Charadriiformes | Europe | TRUE | TRUE | TRUE |
| EPI_ISL_328988 | H13N2 | Charadriiformes | Europe | TRUE | TRUE | TRUE |
| EPI_ISL_328980 | H13N2 | Charadriiformes | Europe | TRUE | TRUE | TRUE |
| EPI_ISL_328984 | H13N2 | Charadriiformes | Europe | TRUE | TRUE | TRUE |
| EPI_ISL_328989 | H13N2 | Charadriiformes | Europe | TRUE | TRUE | TRUE |
| EPI_ISL_328978 | H13N2 | Charadriiformes | Europe | TRUE | TRUE | TRUE |
| EPI_ISL_16919094 | H13N2 | Charadriiformes | North America | TRUE | FALSE | FALSE |
| EPI_ISL_308784 | H13N2 | Charadriiformes | Asia | FALSE | TRUE | TRUE |
| EPI_ISL_16920549 | H13N6 | Charadriiformes | North America | TRUE | FALSE | FALSE |
| EPI_ISL_18059682 | H13N6 | Charadriiformes | Asia | TRUE | TRUE | TRUE |
| EPI_ISL_252213 | H13N6 | Charadriiformes | North America | TRUE | TRUE | TRUE |
| EPI_ISL_252199 | H13N6 | Charadriiformes | North America | TRUE | TRUE | TRUE |
| EPI_ISL_252208 | H13N6 | Charadriiformes | North America | TRUE | TRUE | TRUE |
| EPI_ISL_252198 | H13N6 | Charadriiformes | North America | TRUE | TRUE | TRUE |
| EPI_ISL_252197 | H13N6 | Charadriiformes | North America | TRUE | TRUE | TRUE |
| EPI_ISL_252216 | H13N6 | Charadriiformes | North America | TRUE | TRUE | TRUE |
| EPI_ISL_252207 | H13N6 | Charadriiformes | North America | TRUE | TRUE | TRUE |
| EPI_ISL_252206 | H13N6 | Charadriiformes | North America | TRUE | FALSE | TRUE |
| EPI_ISL_17373343 | H13N6 | Charadriiformes | North America | TRUE | FALSE | FALSE |
| EPI_ISL_4061163 | H13N6 | Charadriiformes | North America | TRUE | FALSE | FALSE |
| EPI_ISL_4056657 | H13N6 | Charadriiformes | North America | TRUE | FALSE | FALSE |
| EPI_ISL_14767321 | H13N6 | Charadriiformes | North America | TRUE | FALSE | FALSE |
| EPI_ISL_9589464 | H13N6 | Charadriiformes | North America | TRUE | TRUE | TRUE |
| EPI_ISL_9589576 | H13N6 | Charadriiformes | North America | TRUE | TRUE | TRUE |
| EPI_ISL_9589331 | H13N6 | Charadriiformes | North America | TRUE | TRUE | TRUE |
| EPI_ISL_9589574 | H13N6 | Charadriiformes | North America | TRUE | TRUE | TRUE |
| EPI_ISL_9589507 | H13N6 | Charadriiformes | North America | TRUE | TRUE | TRUE |
| EPI_ISL_9589438 | H13N6 | Charadriiformes | North America | FALSE | TRUE | TRUE |
| EPI_ISL_9589378 | H13N6 | Charadriiformes | North America | FALSE | TRUE | TRUE |
| EPI_ISL_7622539 | H13N8 | Charadriiformes | Europe | TRUE | TRUE | TRUE |
| EPI_ISL_3319119 | H13N8 | Charadriiformes | Europe | TRUE | TRUE | TRUE |
| EPI_ISL_252219 | H13N8 | Charadriiformes | North America | TRUE | TRUE | TRUE |
| EPI_ISL_252217 | H13N8 | Charadriiformes | North America | TRUE | TRUE | TRUE |
| EPI_ISL_327607 | H13N8 | Charadriiformes | South America | TRUE | TRUE | FALSE |
| EPI_ISL_277382 | H13N8 | Anseriformes | Asia | FALSE | TRUE | TRUE |
| EPI_ISL_308785 | H13N8 | Charadriiformes | Asia | FALSE | TRUE | TRUE |
| EPI_ISL_256303 | H13N8 | environment | Europe | FALSE | TRUE | TRUE |
| EPI_ISL_327624 | H13N8 | Charadriiformes | South America | FALSE | TRUE | FALSE |
| EPI_ISL_381339 | H13N8 | Anseriformes | Asia | FALSE | TRUE | TRUE |
| EPI_ISL_327625 | H13N9 | Charadriiformes | South America | TRUE | TRUE | FALSE |
| EPI_ISL_17978928 | H16N3 | Charadriiformes | Europe | TRUE | TRUE | FALSE |
| EPI_ISL_267201 | H16N3 | Charadriiformes | Europe | TRUE | TRUE | FALSE |
| EPI_ISL_328987 | H16N3 | Charadriiformes | Europe | TRUE | TRUE | TRUE |
| EPI_ISL_376203 | H16N3 | Charadriiformes | Europe | TRUE | TRUE | FALSE |
| EPI_ISL_243597 | H16N3 | Charadriiformes | Europe | TRUE | TRUE | FALSE |
| EPI_ISL_243560 | H16N3 | Charadriiformes | Europe | TRUE | TRUE | FALSE |
| EPI_ISL_243463 | H16N3 | Charadriiformes | Europe | TRUE | TRUE | FALSE |
| EPI_ISL_243545 | H16N3 | Charadriiformes | Europe | TRUE | TRUE | TRUE |
| EPI_ISL_328991 | H16N3 | Charadriiformes | Europe | TRUE | TRUE | FALSE |
| EPI_ISL_328981 | H16N3 | Charadriiformes | Europe | TRUE | TRUE | FALSE |
| EPI_ISL_16920547 | H16N3 | Charadriiformes | North America | TRUE | FALSE | FALSE |
| EPI_ISL_16919118 | H16N3 | Charadriiformes | North America | TRUE | FALSE | FALSE |
| EPI_ISL_399571 | H16N3 | Charadriiformes | Europe | TRUE | TRUE | TRUE |
| EPI_ISL_328974 | H16N3 | Charadriiformes | Europe | FALSE | TRUE | TRUE |
| EPI_ISL_15052970 | H16N3 | Charadriiformes | Asia | FALSE | TRUE | TRUE |
| EPI_ISL_337414 | H16N3 | Charadriiformes | Asia | FALSE | TRUE | TRUE |
| EPI_ISL_277414 | H16N3 | Charadriiformes | North America | FALSE | TRUE | TRUE |
| EPI_ISL_277410 | H16N3 | Charadriiformes | North America | FALSE | TRUE | TRUE |
| EPI_ISL_277416 | H16N3 | Charadriiformes | North America | FALSE | FALSE | TRUE |
| EPI_ISL_277413 | H16N3 | Charadriiformes | North America | FALSE | FALSE | TRUE |
| EPI_ISL_277409 | H16N3 | Charadriiformes | North America | FALSE | FALSE | TRUE |
| EPI_ISL_277408 | H16N3 | Charadriiformes | North America | FALSE | FALSE | TRUE |
| EPI_ISL_277407 | H16N3 | Charadriiformes | North America | FALSE | FALSE | TRUE |
| EPI_ISL_277374 | H16N3 | Charadriiformes | North America | FALSE | FALSE | TRUE |
| EPI_ISL_277415 | H16N3 | Charadriiformes | North America | FALSE | FALSE | TRUE |
| EPI_ISL_277412 | H16N3 | Charadriiformes | North America | FALSE | FALSE | TRUE |
| EPI_ISL_277411 | H16N3 | Charadriiformes | North America | FALSE | FALSE | TRUE |
| EPI_ISL_326959 | H16N3 | Charadriiformes | South America | FALSE | FALSE | TRUE |

## Notes

### Competing Interest Statement

The authors have declared no competing interest.

## References

1. Webster RG, Bean WJ, Gorman OT, Chambers TM, Kawaoka Y. Evolution and ecology of influenza A viruses. Microbiol Rev. 1992 Mar;56(1):152–79.

2. Chen R, Holmes EC. Avian Influenza Virus Exhibits Rapid Evolutionary Dynamics. Molecular Biology and Evolution. 2006 Dec;23(12):2336–41.

3. Hill NJ, Bishop MA, Trovão NS, Ineson KM, Schaefer AL, Puryear WB, et al. Ecological divergence of wild birds drives avian influenza spillover and global spread. Murcia PR, editor. PLoS Pathog. 2022;18(5):e1010062.

4. Lycett SJ, Duchatel F, Digard P. A brief history of bird flu. Phil Trans R Soc B. 2019 24;374(1775):20180257.

5. Rott R. The pathogenic determinant of influenza virus. Veterinary Microbiology. 1992;33(1–4):303–10.

6. Luczo JM, Stambas J, Durr PA, Michalski WP, Bingham J. Molecular pathogenesis of H5 highly pathogenic avian influenza: the role of the haemagglutinin cleavage site motif. Rev Med Virol. 2015;25(6):406–30.

7. Dhingra MS, Artois J, Dellicour S, Lemey P, Dauphin G, Von Dobschuetz S, et al. Geographical and Historical Patterns in the Emergences of Novel Highly Pathogenic Avian Influenza (HPAI) H5 and H7 Viruses in Poultry. Front Vet Sci. 2018;5:84.

8. Duan L, Campitelli L, Fan XH, Leung YHC, Vijaykrishna D, Zhang JX, et al. Characterization of low-pathogenic H5 subtype influenza viruses from Eurasia: Implications for the origin of highly pathogenic H5N1 viruses. J Virol. 2007;81(14):7529–39.

9. Claas EC, Osterhaus AD, Van Beek R, De Jong JC, Rimmelzwaan GF, Senne DA, et al. Human influenza A H5N1 virus related to a highly pathogenic avian influenza virus. The Lancet. 1998;351(9101):472–7.

10. WHO/OIE/FAO H5N1 Evolution Working Group. Continued evolution of highly pathogenic avian influenza A (H5N1): updated nomenclature. Influenza Resp Viruses. 2012;6(1):1–5.

11. Smith GJD, Donis RO, World Health Organization/World Organisation for Animal Health/Food and Agriculture Organization (WHO/OIE/FAO) H5 Evolution Working Group. Nomenclature updates resulting from the evolution of avian influenza A(H5) virus clades 2.1.3.2a, 2.2.1, and 2.3.4 during 2013–2014. Influenza Other Respi Viruses. 2015;9(5):271–6.

12. Lee DH, Bertran K, Kwon JH, Swayne DE. Evolution, global spread, and pathogenicity of highly pathogenic avian influenza H5Nx clade 2.3.4.4. J Vet Sci. 2017;18(S1):269.

13. The Global Consortium for H5N8 and Related Influenza Viruses. Role for migratory wild birds in the global spread of avian influenza H5N8. Science. 2016;354(6309):213–7.

14. Van Den Brand JMA, Verhagen JH, Veldhuis Kroeze EJB, Van De Bildt MWG, Bodewes R, Herfst S, et al. Wild ducks excrete highly pathogenic avian influenza virus H5N8 (2014– 2015) without clinical or pathological evidence of disease. Emerging Microbes & Infections. 2018;7(1):1–10.

15. Xie R, Edwards KM, Wille M, Wei X, Wong SS, Zanin M, et al. The episodic resurgence of highly pathogenic avian influenza H5 virus. Nature;622(7984):810–7.

16. Fusaro A, Zecchin B, Giussani E, Palumbo E, Agüero-García M, Bachofen C, et al. High pathogenic avian influenza A(H5) viruses of clade 2.3.4.4b in Europe—Why trends of virus evolution are more difficult to predict. Virus Evolution. 2024;10(1):veae027.

17. Hagag NM, Yehia N, El-Husseiny MH, Adel A, Shalaby AG, Rabie N, et al. Molecular Epidemiology and Evolutionary Analysis of Avian Influenza A(H5) Viruses Circulating in Egypt, 2019–2021. Viruses. 2022;14(8):1758.

18. Zeng J, Du F, Xiao L, Sun H, Lu L, Lei W, et al. Spatiotemporal genotype replacement of H5N8 avian influenza viruses contributed to H5N1 emergence in 2021/2022 panzootic. Parrish CR, editor. J Virol. 2024;98(3):e01401–23.

19. Olawuyi K, Orole O, Meseko C, Monne I, Shittu I, Bianca Z, et al. Detection of clade 2.3.4.4 highly pathogenic avian influenza H5 viruses in healthy wild birds in the Hadeji-Nguru wetland, Nigeria 2022. Influenza Resp Viruses. 2024;18(2):e13254.

20. Youk S, Torchetti MK, Lantz K, Lenoch JB, Killian ML, Leyson C, et al. H5N1 highly pathogenic avian influenza clade 2.3.4.4b in wild and domestic birds: Introductions into the United States and reassortments, December 2021–April 2022. Virology. 2023;587:109860.

21. Caliendo V, Lewis NS, Pohlmann A, Baillie SR, Banyard AC, Beer M, et al. Transatlantic spread of highly pathogenic avian influenza H5N1 by wild birds from Europe to North America in 2021. Sci Rep. 2022;12(1):11729.

22. Leguia M, Garcia-Glaessner A, Muñoz-Saavedra B, Juarez D, Barrera P, Calvo-Mac C, et al. Highly pathogenic avian influenza A (H5N1) in marine mammals and seabirds in Peru. Nat Commun. 2023;14(1):5489.

23. Jimenez-Bluhm P, Siegers JY, Tan S, Sharp B, Freiden P, Johow M, et al. Detection and phylogenetic analysis of highly pathogenic A/H5N1 avian influenza clade 2.3.4.4b virus in Chile, 2022. Emerging Microbes & Infections. 2023;12(2):2220569.

24. Banyard AC, Bennison A, Byrne AMP, Reid SM, Lynton-Jenkins JG, Mollett B, et al. Detection and spread of high pathogenicity avian influenza virus H5N1 in the Antarctic Region. Nat Commun. 2024;15(1):7433.

25. Signore AV, Giacinti J, Jones MEB, Erdelyan CNG, McLaughlin A, Alkie TN, et al. Spatiotemporal reconstruction of the North American A(H5N1) outbreak reveals successive lineage replacements by descendant reassortants. Sci Adv. 2025;11(28).

26. Agüero M, Monne I, Sánchez A, Zecchin B, Fusaro A, Ruano MJ, et al. Highly pathogenic avian influenza A(H5N1) virus infection in farmed minks, Spain, October 2022. Eurosurveillance. 2023;28(3).

27. Uhart MM, Vanstreels RET, Nelson MI, Olivera V, Campagna J, Zavattieri V, et al. Epidemiological data of an influenza A/H5N1 outbreak in elephant seals in Argentina indicates mammal-to-mammal transmission. Nat Commun. 2024;15(1).

28. Burrough ER, Magstadt DR, Petersen B, Timmermans SJ, Gauger PC, Zhang J, et al. Highly Pathogenic Avian Influenza A(H5N1) Clade 2.3.4.4b Virus Infection in Domestic Dairy Cattle and Cats, United States, 2024. Emerg Infect Dis. 2024;30(7).

29. Koopmans MPG, Barton Behravesh C, Cunningham AA, Adisasmito WB, Almuhairi S, Bilivogui P, et al. The panzootic spread of highly pathogenic avian influenza H5N1 sublineage 2.3.4.4b: a critical appraisal of One Health preparedness and prevention. The Lancet Infectious Diseases. 2024;S1473309924004389.

30. Peacock T, Moncla L, Dudas G, VanInsberghe D, Sukhova K, Lloyd-Smith JO, et al. The global H5N1 influenza panzootic in mammals. Nature. 2024; Available from: https://www.nature.com/articles/s41586-024-08054-z

31. Klaassen M, Wille M. The plight and role of wild birds in the current bird flu panzootic. Nat Ecol Evol. 2023;7(10):1541–2.

32. Bedford T, Riley S, Barr IG, Broor S, Chadha M, Cox NJ, et al. Global circulation patterns of seasonal influenza viruses vary with antigenic drift. Nature. 2015;523(7559):217–20.

33. Himsworth CG, Caleta JM, Jassem AN, Yang KC, Zlosnik JEA, Tyson JR, et al. Highly Pathogenic Avian Influenza A(H5N1) in Wild Birds and a Human, British Columbia, Canada, 2024. Emerg Infect Dis. 2025;31(6).

34. Peacock TP, Sheppard CM, Staller E, Barclay WS. Host determinants of influenza RNA synthesis. Annu Rev Virol. 2019;6(1):215–33.

35. Long JS, Idoko-Akoh A, Mistry B, Goldhill D, Staller E, Schreyer J, et al. Species specific differences in use of ANP32 proteins by influenza A virus. eLife. 2019;8:e45066.

36. Staller E, Sheppard CM, Neasham PJ, Mistry B, Peacock TP, Goldhill DH, et al. ANP32 Proteins Are Essential for Influenza Virus Replication in Human Cells. García-Sastre A, editor. J Virol. 2019;93(17):e00217–19.

37. Ji Z xing, Wang X quan, Liu X fan. NS1: A Key Protein in the “Game” Between Influenza A Virus and Host in Innate Immunity. Front Cell Infect Microbiol. 2021 Jul 13;11:670177.

38. Jagger BW, Wise HM, Kash JC, Walters KA, Wills NM, Xiao YL, et al. An overlapping protein-coding region in influenza A virus segment 3 modulates the host response. Science. 2012;337(6091):199–204.

39. Shu Y, McCauley J. GISAID: Global initiative on sharing all influenza data – from vision to reality. Eurosurveillance. 2017;22(13).

40. Altschul SF, Gish W, Miller W, Myers EW, Lipman DJ. Basic local alignment search tool. Journal of Molecular Biology. 1990;215(3):403–10.

41. Verhagen JH, Poen M, Stallknecht DE, Van Der Vliet S, Lexmond P, Sreevatsan S, et al. Phylogeography and Antigenic Diversity of Low-Pathogenic Avian Influenza H13 and H16 Viruses. Parrish CR, editor. J Virol. 2020;94(13):e00537–20.

42. Byrne AMP, James J, Mollett BC, Meyer SM, Lewis T, Czepiel M, et al. Investigating the Genetic Diversity of H5 Avian Influenza Viruses in the United Kingdom from 2020–2022. Deng T, editor. Microbiol Spectr. 2023;11(4):e04776–22.

43. Gamża A, Lycett S, Sanchez A, Kao R. Using sequence data to study spatial scales of interactions driving spread of Highly Pathogenic Avian Influenza in Great Britain. arXiv; 2024. Available from: https://arxiv.org/abs/2411.10424

44. Katoh K, Standley DM. MAFFT Multiple Sequence Alignment Software Version 7: Improvements in Performance and Usability. Molecular Biology and Evolution. 2013;30(4):772–80.

45. Tavaré, S. Some probabilistic and statistical problems in the analysis of DNA sequences. Lectures on Mathematics in the Life Sciences. 1986;17:57–86.

46. Price MN, Dehal PS, Arkin AP. FastTree: Computing Large Minimum Evolution Trees with Profiles instead of a Distance Matrix. Molecular Biology and Evolution. 2009;26(7):1641–50.

47. R Core Team. R: A language and environment for statistical computing. [Internet]. Vienna, Austria: R Foundation for Statistical Computing; 2023. Available from: https://www.R-project.org/

48. Keck F. Handling biological sequences in R with the bioseq package. Mahon A, editor. Methods Ecol Evol. 2020;11(12):1728–32.

49. Suchard MA, Lemey P, Baele G, Ayres DL, Drummond AJ, Rambaut A. Bayesian phylogenetic and phylodynamic data integration using BEAST 1.10. Virus Evolution. 2018;4(1).

50. Shapiro B, Rambaut A, Drummond AJ. Choosing Appropriate Substitution Models for the Phylogenetic Analysis of Protein-Coding Sequences. Molecular Biology and Evolution. 2006;23(1):7–9.

51. Hasegawa M, Kishino H, Yano T. Dating of the human-ape splitting by a molecular clock of mitochondrial DNA. J Mol Evol. 1985;22(2):160–74.

52. Lanave C, Preparata G, Sacone C, Serio G. A new method for calculating evolutionary substitution rates. J Mol Evol. 1984;20(1):86–93.

53. Drummond AJ, Ho SYW, Phillips MJ, Rambaut A. Relaxed Phylogenetics and Dating with Confidence. Penny D, editor. PLoS Biol. 2006;4(5):e88.

54. Gill MS, Lemey P, Faria NR, Rambaut A, Shapiro B, Suchard MA. Improving Bayesian Population Dynamics Inference: A Coalescent-Based Model for Multiple Loci. Molecular Biology and Evolution. 2013;30(3):713–24.

55. Yu G, Smith DK, Zhu H, Guan Y, Lam TT. GGTREE : an R package for visualization and annotation of phylogenetic trees with their covariates and other associated data. McInerny G, editor. Methods Ecol Evol. 2017;8(1):28–36.

56. Lemey P, Rambaut A, Drummond AJ, Suchard MA. Bayesian Phylogeography Finds Its Roots. PLoS Comput Biol. 2009;5(9):e1000520.

57. Lemey P, Rambaut A, Welch JJ, Suchard MA. Phylogeography Takes a Relaxed Random Walk in Continuous Space and Time. Molecular Biology and Evolution. 2010;27(8):1877–85.

58. McIlroy D, Brownrigg R, Minka TP, Deckmyn A. mapproj: Map Projections.

59. Harvey WT, Mulatti P, Fusaro A, Scolamacchia F, Zecchin B, Monne I, et al. Spatiotemporal reconstruction and transmission dynamics during the 2016–17 H5N8 highly pathogenic avian influenza epidemic in Italy. Transbound Emerg Dis. 2021;68(1):37–50.

60. Neher RA, Russell CA, Shraiman BI. Predicting evolution from the shape of genealogical trees. eLife. 2014;3:e03568.

61. Barido-Sottani J, Vaughan TG, Stadler T. A Multitype Birth–Death Model for Bayesian Inference of Lineage-Specific Birth and Death Rates. Systematic Biology. 2020;69(5):973–86.

62. Bouckaert R, Heled J, Kühnert D, Vaughan T, Wu CH, Xie D, et al. BEAST 2: A Software Platform for Bayesian Evolutionary Analysis. PLoS Comput Biol. 2014;10(4):e1003537.

63. Kühnert D, Stadler T, Vaughan TG, Drummond AJ. Phylodynamics with Migration: A Computational Framework to Quantify Population Structure from Genomic Data. Mol Biol Evol. 2016;33(8):2102–16.

64. Esnault E, Bonsergent C, Larcher T, Bed’hom B, Vautherot JF, Delaleu B, et al. A novel chicken lung epithelial cell line: Characterization and response to low pathogenicity avian influenza virus. Virus Research. 2011;159(1):32–42.

65. Sheppard CM, Goldhill DH, Swann OC, Staller E, Penn R, Platt OK, et al. An Influenza A virus can evolve to use human ANP32E through altering polymerase dimerization. Nat Commun. 2023;14(1):6135.

66. de Wit E, Spronken MIJ, Bestebroer TM, Rimmelzwaan GF, Osterhaus ADME, Fouchier RAM. Efficient generation and growth of influenza virus A/PR/8/34 from eight cDNA fragments. Virus Research. 2004;103(1–2):155–61.

67. Moncorgé O, Long JS, Cauldwell AV, Zhou H, Lycett SJ, Barclay WS. Investigation of Influenza Virus Polymerase Activity in Pig Cells. J Virol. 2013;87(1):384–94.

